# Toward an understanding of the relation between gene regulation and 3D genome organization

**DOI:** 10.1101/2020.01.13.903872

**Authors:** Hao Tian, Ying Yang, Sirui Liu, Hui Quan, Yi Qin Gao

**Affiliations:** Beijing National Laboratory for Molecular Sciences, College of Chemistry and Molecular Engineering, Peking University, Beijing 100871, China; Biomedical Pioneering Innovation Center (BIOPIC), Peking University, Beijing 100871, China; Beijing Advanced Innovation Center for Genomics (ICG), Peking University, Beijing 100871, China

## Abstract

The development and usage of chromosome conformation capture technologies have provided great details on 3D genome organization and provide great opportunities to understand how gene regulation is affected by the 3D chromatin structure. Previously, we identified two types of sequence domains, CGI forest and CGI prairie, which tend to segregate spatially, but to different extent in different tissues/cell states. To further quantify the association of domain segregation with gene regulation and differentiation, we analyzed in this study the distribution of genes of different tissue specificities along the linear genome, and found that the distribution patterns are distinctly different in forests and prairies. The tissue-specific genes (TSGs) are significantly enriched in the latter but not in the former and genes of similar expression profiles among different cell types (co-activation/repression) also tend to cluster in specific prairies. We then analyzed the correlation between gene expression and the spatial contact revealed in Hi-C measurement. Tissue-specific forest-prairie contact formation was found to correlate with the regulation of the TSGs, in particular those in the prairie domains, pointing to the important role gene positioning, in the linear DNA sequence as well as in 3D chromatin structure, plays in gene regulatory network formation.

## Introduction

Over the last decade, with the development of chromosome conformation capture technologies, such as Hi-C (*1*) and ChIA-PET (*2*), people have gained much insight into the high order chromatin structure and a series of intriguing discoveries (for example, compartment (*1*), TAD (*3*) and chromatin loop (*4*)) have been made. In terms of the relation between structural organization and gene regulation, chromatin loops, frequently anchored by CCCTC-binding factor (CTCF) and cohesin complex, can link promoter regions and distal regulatory elements such as enhancers to ensure normal transcription (*4*). Such spatial interactions are often restricted into individual specific topological domains, named insulated neighbors that can be seen as a mechanistic basis of TADs (*5*, *6*). TAD boundaries are often thought to be robust among different cell types (*3*) and the relationship between TAD formation and gene regulation has caused much research interest. For example, the disruption of specific TAD boundaries or insulated structures and consequently unexpected enhancer-promoter interactions can induce aberrant gene expression, especially that of disease-related genes (*7*–*9*). Surprisingly, the global loss of TAD structure via depletion of cohesin leads to significant changes of expression level of only about 1000 genes (*10*), indicating a limited role of TAD structure in regulating gene transcription. Moreover, multiplex promoter-centered chromatin interactions such as promoter-promoter interactions of different genes have been widely investigated using ChIA-PET targeting on RNA polymerase II (RNAPII) and are thought to play a vital role in transcription regulation (*11*). A-B compartmentalization has been referred to efficiently participate in gene regulation possibly owing to the local enrichment of transcription-associated factors (*12*) and compartment shifts have been found to coincide with the activation or repression of specific genes (*13*, *14*). However, a systematic understanding of regulation of tissue-specific genes (TSGs) from the perspective of chromatin organization is still in need.

Besides expression levels of individual genes, the correlation between the expression levels of different genes in connection with their positions in 3D space provides additional information on biological function regulation. Gene networks, constructed mainly based on pairwise gene co-regulation patterns, summarize the expression correlation between genes and can serve as a powerful tool to identify regulatory modules (*15*–*17*). Furthermore, in recent years, phase separation models, describing a phenomenon in which specific proteins, such as transcription factors (TFs), co-activators, and RNAPⅡ, can accumulate and self-organize into condensates, have been proposed to contribute to the formation of 3D chromatin structure as well as gene regulation (*18*–*20*). Such a model is accordant with observations on the co-location of TF-binding sites and corresponding genes (*21*, *22*). Despite of all these efforts, it remains largely unexplored about the interplay between gene pair expression coordination and high order chromatin structure. It is thus interesting to examine whether spatial contact also plays a role in gene co-regulation network formation.

In this paper, we investigate in a systematic way how gene expression is related to chromatin structure. One of the important factors affecting gene expression is the compartmentalization (*12*–*14*), which was found to be strongly affected by the DNA sequence (*23*). Based on the uneven distribution of CpG islands (CGIs), the whole genome was divided into two types of domains, named CGI forest (F) and CGI prairie (P), respectively. The former is enriched in CGI, possesses high gene (especially housekeeping gene) density and active histone marks, whereas the latter is characterized by low CGI and gene densities, and strong signals of repressive histone marks. More importantly, different extent of domain segregation between forests and prairies was observed in different cell types. This domain segregation correlates with the establishment of cell identity, providing a description on how compartmentalization differs among different cell types.

Here, we first explore the distribution of genes of different tissue specificities in the linear genome, in particular their different distribution patterns in forest and prairie domains. Prairies are found to be more likely than forests to contain genes of high tissue specificity. We then identified functional modules for different cell types, which are individual prairie domains that significantly encompass large numbers of genes that are specifically highly expressed in a cell-specific way. Next, we found that genes of higher tissue specificity in prairies but not forests are prone to be characterized by lower gene body CpG density. Based on earlier findings on the negative correlation between GC content (or gene body CpG density) and repressive histone mark H3K9me3 (*24*, *25*) and our analysis on the positive association between gene body CpG density and compartment index, we inferred that prairie genes of higher tissue specificities tend to reside in repressed (heterochromatin-like) environment except when they are highly expressed in a tissue (cell state)-specific manner. Furthermore, how TSGs are activated in corresponding cells from the perspective of spatial interactions between forest and prairie domains is discussed. We propose that the achievement of cell identity is realized with the help of spatial contacts formed between prairie and forest genes, through the assistant of TFs. To further explore the association between gene pair expression coordination and genome organization, we constructed gene co-regulation networks based on expression correlation calculation, and found that in these networks genes tend to correlate with other genes in the same compartment. We then examined the spatial interaction patterns co-regulated genes formed through constructing gene networks based on a combined usage of Hi-C (*26*) and RNA-seq (*27*) data, from which we observed that the F-P gene expression correlation patterns are indeed coupled to the overall chromatin structure features. Finally, the quantitative assessment of the relation between gene co-regulation and high order chromatin structure is performed and how spatial contacts among genes affect their expression coordination is discussed.

## Materials and methods

### Definition of gene tissue specificity

The tissue specificity of gene *i* in tissue *t* is defined as

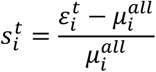

where 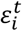 and 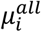 are the mean expression level of gene *i* in tissue *t* and all tissue types included in the calculation (see Supplementary File 1), respectively. We consider gene *i* to be specific to tissue *t* if 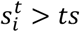. Different values of the cutoff *ts* were tested and the results are robust to this parameter. Without specific notation, the results we presented were obtained using a *ts* = 2.

### Saturation curve of the number of relevant TSGs at individual forest/prairie domain level

To better describe the distribution of relevant TSGs at individual domain level, the vector composed of the number (non-zero) of forest/prairie TSGs related to one cell (e.g., liver) in specific individual domain was first sorted in a descending order and then each element, e.g., the *i*^th^, was converted to the ratio between the summation of the first *i* numbers and the total number of forest/ prairie TSGs belonging to one cell (e.g., liver). For a given value of y, the corresponding x value was normalized to the ratio between *i* and the total number of forest (766) /prairie (801) domains.

### Definition of segregation ratio

Segregation ratio of a prairie domain *i,* 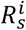, is defined as (*23*)

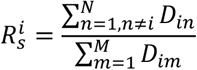

where *N* (n∈P) and *M* (m∈F) are the collection of all prairie and forest domains, respectively. *D*_*ij*_ is the summation of normalized Hi-C contact probability between domains *i* and *j*. Without specific notation, the Hi-C contact matrices were normalized using ICE (iterative correction and eigenvector decomposition) method (*28*). A high value of 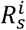 implies a high probability of the prairie domain *i* to be spatially in contact with prairie regions.

### Calculation of compartment index

The compartment index of bin *i*, *CI*_*i*_, is calculated following our previous work (*29*) as

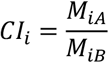

where *M*_*iA*_ and *M*_*iB*_ are the mean spatial contact probabilities of bin *i* with compartments A and B, respectively. The identification of compartment A/B follows our previous work with slight modifications(*23*): the entire Hi-C matrix was disassembled into two parts, corresponding to p and q arms, and the eigenvalue decomposition was done within these two arms separately. A high value of *CI*_*i*_ suggests that bin *i* locates in a more compartment A environment.

### Calculation of forest index

Forest index of a 40-kb bin *i* is defined as (*23*),

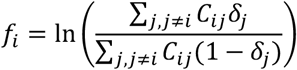

where *C*_*ij*_ is the contact probability between bins *i* and *j*. *δ*_*j*_ equals to 1 if the 40-kb bin *j* locates in a forest, and 0 if it is in a prairie domain. Therefore, a larger value of *f*_*i*_ indicates that the 40-kb bin is spatially surrounded by a higher population of forests.

### Construction of gene networks

Gene networks displayed in this study were constructed based on a combined usage of Hi-C (*26*) and RNA-seq (*27*) data. Two genes connected by an edge in the network not only possess a high co-regulation level, but are also in strong spatial proximity in terms of Hi-C contact.

Two genes *i* and *j* are considered to be under co-regulation, if the Pearson correlation coefficient calculated from 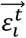 and 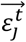 exceeds the 99.5^th^ percentile of all correlations {*P*_*ij*_} (*i*, *j* ∈ {1,2, …, *N*} and *i* < *j*, *N* is the number of genes in the dataset we considered, based on such criterion we constructed gene co-regulation network).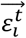 and 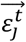 are the expression vectors for genes *i* and *j* in tissue *t*, respectively. Each element of the expression vector represents the expression level of one sample belonging to tissue *t*. For instance, if tissue *t* has *n*_*t*_ samples in the corresponding RNA-seq data, the size of 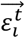 is 1 × *n*_*t*_.

The Hi-C contact matrices are of 40-kb resolution and the co-regulated gene pairs *i* and *j* are therefore projected to 40-kb bins, denoted as *a* and *b*, respectively. The genomic distance between *a* and *b* is denoted as *d*_*ab*_. Given that the contact probability decays quickly with the genomic separation, we use relative values to evaluate the strength of gene pair spatial interaction. Namely, genes *i* and *j* are regarded in strong spatial contact in tissue *t* if the normalized Hi-C contact probability of bins *a* and *b*, 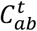,exceeds the 75^th^ percentile (90^th^ percentile was used for robustness test and similar results were acquired) of 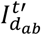 that was defined as

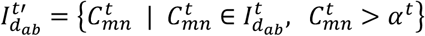

where 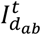 is the vector composed of spatial contacts between two bins, the genomic distance of which is equal to *d*_*ab*_. *α*^*t*^ is defined as

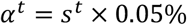

where *s*^*t*^ is the summation of one row in tissue *t*’s normalized Hi-C contact matrix. Such value was used to align the Hi-C data of different cells to similar scale (The concrete values and results can be found in **Table S1**).

### Identification of gene functions

The website https://www.genecards.org was used for the analysis of gene functions.

### Definition of Hi-C rank

The Hi-C rank *r*_*i*,*j*_, between two genes *i* and *j*, is defined as

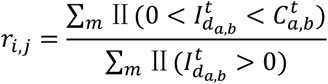

with 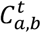 and 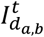 defined earlier. As the genomic distance increases, *N*_*n*_⁄*Z*_*n*_ decays exponentially (**Figure S1**), where *N*_*n*_ and *Z*_*n*_ represent the number of non-zero and zero elements in the Hi-C matrix at the given genomic distance. Thus, we retained only non-zero elements and calculated the rank within them.

## Results

### Forest and prairie gene features

As mentioned earlier, we divided the whole genome into two types of domains, forests and prairies, solely based on DNA sequence properties. Different levels of forest-prairie segregation were observed in different cells, showing a strong sequence dependence in chromatin compartmentalization. In order to further clarify the underlying biological significances of domain segregation (e.g. the activation of specific genes), we firstly investigated the distribution of genes of different tissue specificities as well as the enrichment of TSGs between the two types of linear domains. It can be seen from **Table 1** that TSGs relevant to one certain cell type tend to be enriched in prairies rather than forests in most tissues we considered (the tissues we mainly analyzed in this study can be seen in Supplementary text), except for those possessing very small numbers of TSGs, such as aorta, left ventricle and ovary. The distribution of gene tissue specificity (see methods) in these two kinds of domains is also distinctly different (**Figure 1A and S2**). The proportion of genes of high (positive) and extremely negative tissue specificity (a gene possessing a negative tissue specificity is one that highly expressed in some other tissues rather than the reference tissue) is larger in prairies than in forests, corresponding to the enrichment of TSGs in the relevant and other cell types, respectively. In contrast, forests are abundant in genes, the tissue specificity distribution of which is centered at zero (not tissue-specific).

**Figure 1.**
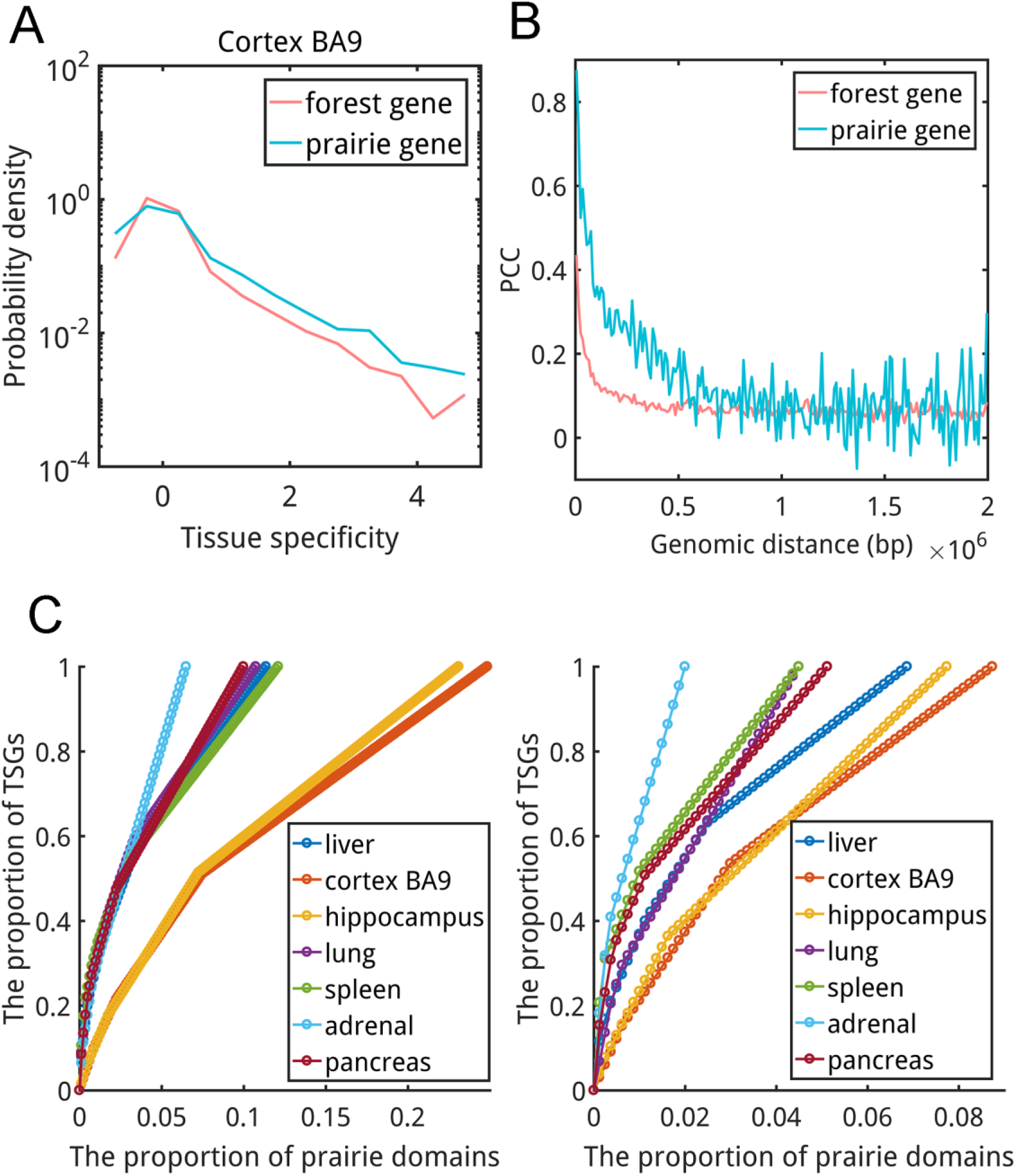
Forest and prairie gene features. (**A**) The distribution profile of gene tissue specificities for forests and prairies in cortex BA9. (**B**) Pearson correlation coefficient (PCC) calculated between the tissue specificities (among different cell types) of two forest or prairie genes within the same domain. (**C**) Distribution of TSGs (left, 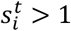; right, 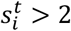) in prairies (see methods).

**Table 1.** TSG distribution in forest and prairie domains.

|  | Liver | Cortex | Hippo | Lung | LV | Spleen | Ovary | Adrenal | Aorta | Panc |
| --- | --- | --- | --- | --- | --- | --- | --- | --- | --- | --- |
| F_t | 217 | 193 | 169 | 67 | 40 | 106 | 43 | 55 | 10 | 128 |
| F_nt | 14853 | 14877 | 14901 | 15003 | 15030 | 14964 | 15027 | 15015 | 15060 | 14942 |
| P_t | 95 | 99 | 77 | 44 | 8 | 58 | 11 | 22 | 4 | 65 |
| P_nt | 3248 | 3244 | 3266 | 3299 | 3335 | 3285 | 3332 | 3321 | 3339 | 3278 |
| *P*-val | 1.1e-7 | 9.7e-11 | 6.4e-7 | 9.5e-8 | - | 1.3e-7 | - | 0.025 | - | 2.4e-7 |

\*F\_t, F\_nt, P\_t and P\_nt represent the number of forest TSG, forest non-TSG, prairie TSG and prairie non-TSG in corresponding tissue, respectively. Fisher's exact test was performed and "-" indicates the corresponding p-value is larger than 0.05. Hippo=hippocampus, LV=left ventricle, Panc=pancreas.

It is known that genes of related functions can cluster along the linear genome (*30*, *31*). To quantify the clustering of function-related genes within individual forest or prairie domains, we calculated the Pearson correlation coefficient (PCC) of two genes’ tissue specificity among different tissues. It was found that forest genes are less correlated than prairies (**Figure 1B**, each data point represents the median value of PCC derived from a group of gene pairs, the genomic distance of which lies within one 10-kb-long range, e.g., [0,1000bp),[10000bp,20000bp)). This result indicates that when the cell type changes, forest genes tend to express independently to their neighboring genes within the same forest domain, whereas prairie genes display a much stronger tendency of co-activation/repression with its nearby genes. Thus, prairie genes of similar expression profiles (and probably related biological functions) are more likely to cluster in the specific genome region, readily positioned for synergistic regulation. A number of prairie domains are identified to significantly enrich TSGs of similar functions and we term these domains as functional modules (**Figure S3A**). A functional module is defined based on the condition that the number of relevant TSGs (e.g., liver TSGs) in this prairie domain is equal to or even greater than the maximum number of related TSGs among forest domains, although forests possess the great majority (78.5%) and thus a much higher density of genes. For instance, 12 out of 27 genes (*P*-val < 10^−18^ by Fisher’s exact test) residing in the 31^st^ prairie domain (PD31) in chr19 (genomic location: 54711241-55450974) are specific to spleen; 24 genes in this domain are related to immune function, so this prairie domain is heavily involved in immune response and maintenance of the normal physiology of immune system. Functional modules were also identified for liver, adrenal and pancreas (**Figure S3A and Table S2**). However, although encompassing considerable numbers of related TSGs, such prairie domains are not found for cortex BA9 and hippocampus (**Figure S3A)**. We want to note here that if a less strict criterion (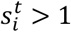, instead of 2) in identifying TSGs is adopted, one functional module (**Figure S3B**) is also identified for lung, which is again PD31, indicating the importance of immune related gene expressions in lung. In contrast, no functional module can be identified in cortex BA9 and hippocampus even with 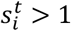 (**Figure S3B**). Our previous analysis revealed that unlike liver and spleen, which are proliferating cells and displaying strong F-P domain separation, especially long-range P-P aggregation, cortex BA9 exhibits a strong mixing between the forest and prairie domains (*23*). The lack of brain-related functional module and the largely uniform distribution of brain-related genes in the genome sequence (**Figure 1C, S3 and S4**) appear to associate with such a chromatin 3-D structural feature, which will be discussed below.

Furthermore, the forest and prairie domains are shown to have a different correlation between gene tissue specificity and gene body CpG density (**Figure 2A**). For prairie but not forest genes, as their tissue specificity increases, the gene body CpG density decreases. The CpG density of prairie housekeeping genes (HKGs) is generally higher than that of prairie TSGs. Several earlier studies have revealed that the repressive histone mark H3K9me3 is strongly enriched in genes characterized by low CpG density in gene body (*24*) and the H3K9me3 density negatively correlates with GC content (*25*). Moreover, we observed a positive correlation between compartment index (see methods) and gene body CpG density (**Figure 2B, S5 and S6**). These results indicate that prairie genes possessing higher tissue specificities tend to be CpG poor in gene body and reside in compartment B unless they are actively transcribed in their corresponding tissue, presumably with the help of specific TFs. On the other hand, prairie HKGs are on average CpG richer than prairie TSGs in gene body, although their transcriptional level is still lower than their counterparts in forests (**Figure S7**).

**Figure 2.**
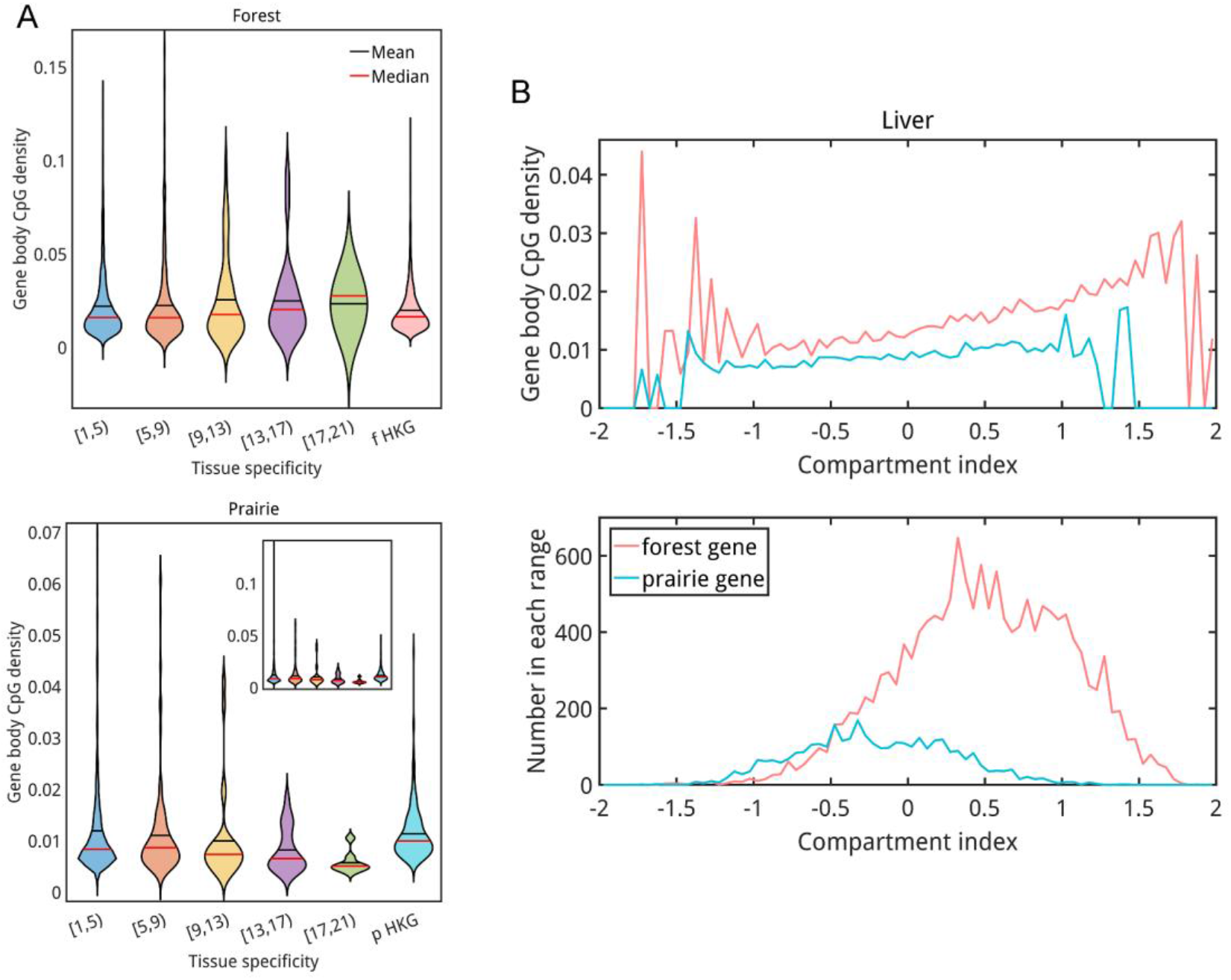
(**A**) The relation between gene tissue specificity and gene body CpG density. The gene tissue specificity is its maximum value among all tissues (except for testis, which contains a very large number of TSGs, see Supplementary file 2). (**B**) The relation between compartment index and gene body CpG density. Upper, the median value of gene body CpG density, the compartment indices of corresponding genes lie within one 0.05-long range, e.g., [−2,−1.95). Down, the number of genes in each range.

### The regulation of genes, especially those in prairies, is closely linked to 3D chromatin structure

It is thus interesting to understand how prairie TSGs, especially those characterized by very low CpG densities in gene body, become highly transcribed in their corresponding tissues (cells). Notably, we found that unlike forest TSGs, the majority of prairie TSGs still reside in compartment B (the corresponding eigenvector is smaller than zero, see methods) even in cells where they are highly transcribed (**Figure 3A**). To examine whether the activation of prairie TSGs can be related to chromatin structural organization, we used forest index (*f*_*i*_, see methods) to evaluate the local spatial environment of prairie TSGs. A high value of *f*_*i*_ implies that a gene is embedded in a spatial environment rich in forest domains. Our analysis revealed that the forest indices of prairie TSGs in the relevant tissue are generally more positive than those of the same genes when they are in other tissues (**Figure 3B and S9A**, the value behind each symbol represents the median forest index difference between related cell (illustrated in legend) and control cell (labeled in x-axis) for related cell’s prairie TSGs), in agreement with the changes of compartment index (**Figure S8A and S9C**). Furthermore, changing from one cell type to another, the changes of forest and compartment indices of prairie TSGs specific to the latter are generally more positive (**Figure 3C, S8B, S8C, S9B and S9D**) than those specific to the other cells. These observations indicate that the TSGs of a given tissue are in a more forest-surrounded and more active 3D structural environment in that particular tissue than in other tissues. Besides, as previously noted, the TSGs are enriched in prairie domains compared to forests and genes of similar expression behaviors (co-activation/repression) are more likely to cluster in specific prairie regions. In this sense, these particular prairie domains can be considered as specified warehouses of TSGs. Collective activation or repression of these genes can then be achieved through concerted and tissue (cell state)-specific 3D chromatin reorganization of these prairie domains, by specifically intermingling with or separating from forest domains.

**Figure 3.**
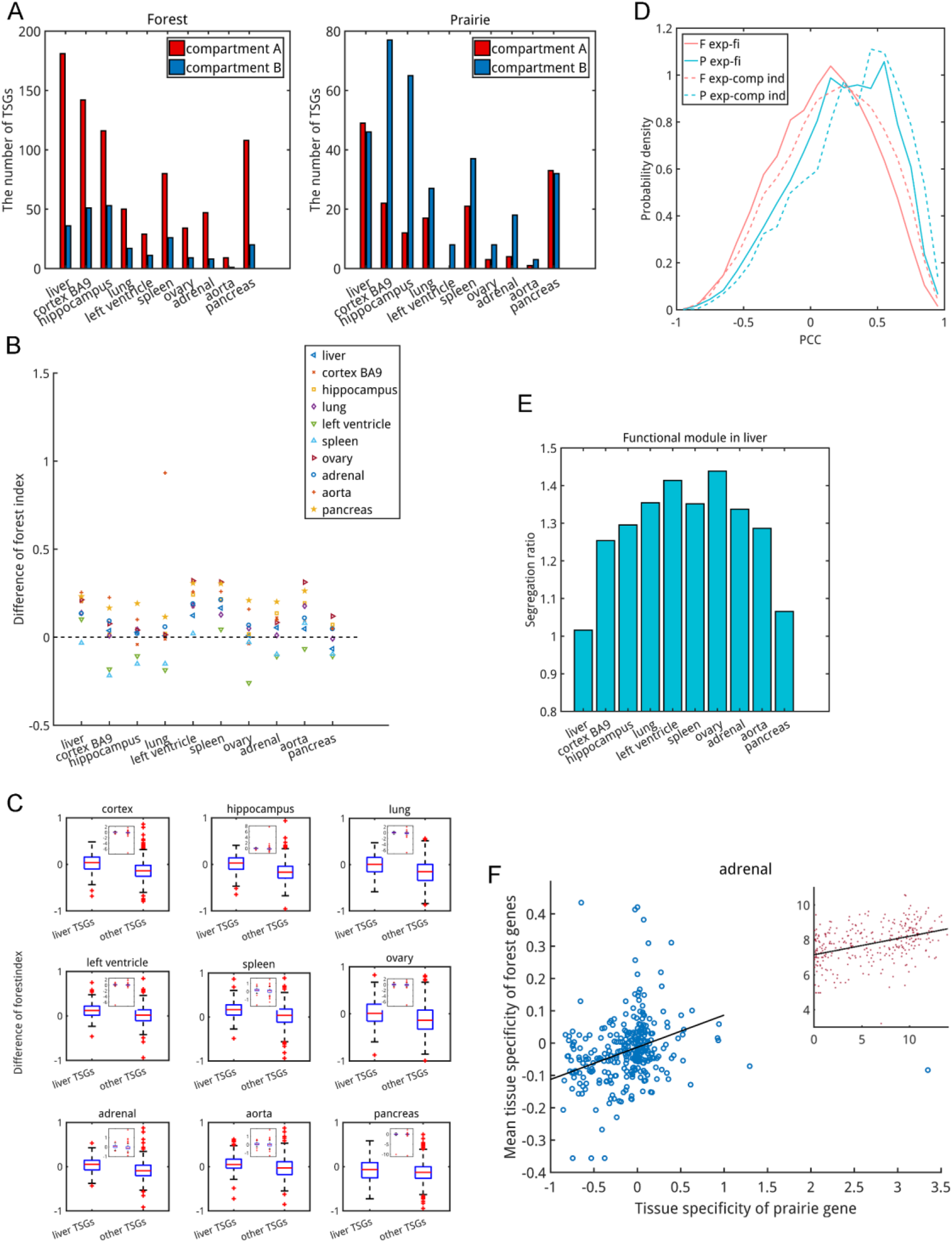
Prairie gene expression regulation and genome organization. (**A**) The number of TSGs residing in compartment A (eigenvector > 0, see methods) or B (eigenvector < 0). (**B**) Forest index changes of prairie TSGs (e.g., liver prairie TSGs) from other cells (e.g. spleen) to the relevant cell (e.g. liver). (**C**) Comparison of forest index changes between relevant TSGs and TSGs specific to other cells. Liver was used as illustration. (**D**) Pearson correlation coefficient (PCC) between expression level and forest/compartment index for forest and prairie genes. (**E**) The segregation ratio of liver functional module in ten cells, since these ten cells were performed both Hi-C and RNA-seq experiments and therefore were mainly used for analysis in this study (see Supplementary text). (**F**) Scatter plot for tissue specificity (expression level, inner figure) of prairie genes and the averaged tissue specificity (expression level) of forest genes in strong contact.

The above phenomena, in which specific spatial interactions with forest domains are coupled with prairie gene regulation, were also observed in forest (**Figure S10 and S11**). However, one can see that the compartment index changes of forest TSGs are generally less significant than prairie TSGs in transcription activation (**Figure S12 and S13**). Furthermore, we calculated the Pearson correlation coefficient between gene expression levels and forest/compartment indices. One can see from **Figure 3D** that the values for forest genes tend to be more negative than those for prairie genes. Therefore, the activation of forest genes depends less on the specific spatial interactions (i.e., movement to forest and compartment A environment) compared to prairie genes, suggesting the importance of other mechanisms other than compartmentalization in forest gene expression regulation. Interestingly, we found that forest genes showing strong negative correlations between their gene expression level and forest index tend to be brain-related (**Figure S14 and S15**). This observation, together with the broader and more uniform distribution of brain prairie TSGs in the genome sequence that need to intermingle with forest regions for activation, is accordant with the cortex BA9 chromatin structure which shows strong intermingling between forest and prairie domains (*23*).

The tissue-specific functional modules defined earlier also show tissue-specific enhanced contacts with forest domains. We calculated the segregation ratio *R*_*s*_ (see methods) to quantify the segregated state (a prairie with other prairies) of these prairie domains. The functional modules in liver, spleen and adrenal all showed the nearly smallest *R*_*s*_ values in the corresponding tissues (**Figure 3E and S16**), indicating the expected F-P intermingling of these domains is consistent with tissue-specific active transcription. Overall, the tissue-specific genome organization is consistent with expression level of genes residing in these domains. The functional module of pancreas was not included in this calculation as the pancreas TSG (amylase gene) cluster locates in a very small, variable and dynamic region (*32*), in which the Hi-C contact information is missing.

As a whole, the aforementioned results are consistent with prairie TSGs entering a more active environment surrounded by a higher population of forests for activation in their corresponding tissues, when compared to the environment of these same genes but in other tissues. In addition, a positive (although weak) correlation was also observed between the tissue specificity of prairie genes and the mean tissue specificity of its highly contacted forest genes (**Figure 3F, S17 and S18**). Such a result indicate that prairie genes of high tissue specificities may tend to contact with the genes of related functions residing in forests. A positive correlation also exists between gene expression levels of prairie genes and its highly contacted forest genes (**Figure 3F inner, S19 and S20**).

### The interplay between gene pair expression coordination and genome organization

Given that the activation and repression of genes are largely associated with genome organization, we then asked whether gene pair expression coordination is also related to high order chromatin structure. To address this question, we first constructed gene networks based on their expression correlation coefficients (see methods). To quantify the effect of compartmentalization on gene expression correlation, we then calculated the average number of compartment X(X=A or B) genes connected to genes in compartment Y(Y=A or B), N_Y-X_, in the co-regulation networks. Interestingly, genes tend to correlate with other genes residing in the same other than different compartment type, since N_B-A_ and N_A-B_ are always lower than N_A-A_ and N_B-B_, respectively, with the exception of cortex BA9, for which the N_B-A_ is larger than N_A-A_, implying the prominent behaviors of compartment B genes in mediating brain cell activity (**Figure 4A, S21**). Such a result strongly suggests a non-negligible association between compartmentalization and gene pair co-regulation. To further explore the spatial interactions of co-regulated gene pairs, we next constructed a new type of gene network, the edges of which connect genes that are not only highly correlated but also are in close spatial proximity (see methods). Two cell samples, liver and cortex BA9, were selected to illustrate the results as their F-P domain segregation patterns are significantly different as shown in our previous work (*23*). The former displays a strong separation between forest and prairie domains and an aggregation of prairie domains distant on the linear genome, whereas the latter shows a strong intermingling between forest and prairie domains. All genes of chr1 were used to construct the gene network. The results revealed that such networks in liver and cortex BA9 are mainly composed of two independent sub-networks (**Figure S22 and S23**). The gene tissue specificity distributions of these two sub-networks are distinctly different (**Figure 4C and S24**). One of them is directly associated with cell identity and the other, of more generic functions, indicating that the 3D chromatin structure and spatial contacts among genes may play an important role in the formation of tissue-specific gene network and the establishment of tissue specificity. In addition, the pattern of interactions between forest and prairie genes in the tissue-specific sub-network does reflect the overall segregation of these types of domains in that particular tissue (**Figure 4B**). In liver, the network core is mainly composed of forest genes and prairie genes occupy only the marginal positions, whereas the cortex BA9 network displays a strong intermingling of forest and prairie genes (the non-tissue-specific sub-network in liver also displays F-P separated state whereas in cortex BA9 the scale of such network is very small, **Figure S22 and S23**). Since unlike the standard gene expression correlation network, the network presented here also includes spatial information and can be used to obtain information on the co-regulatory mechanism (such as the binding of TFs and spatial clustering) of the genes involved. For example, in liver network, *CA14* (a forest gene) is found to connect with six genes, of which five (*HFE2*, *ANXA9*, *SELENBP1*, *PKLR*, *RHBG*) reside in forest domains and one (*FMO3*) in prairie. In liver, *HFE2*, *PKLR*, *RHBG*, *ANXA9* and *FMO3* are highly expressed and the promoter regions of all these seven genes bind HNF4A, a liver-specific TF, RXRA and YY1 (*33*), strongly suggesting that the co-localization of these genes, with the help of TF binding, plays a role in the formation of tissue-specific gene regulation network and their co-activation in liver. Hence, like in the phase separation model, the cell–specific formation of such clusters as well as spatial contacts between forest and prairie genes may render an efficient regulatory mechanism in the gene transcription, especially for prairie genes, and the subsequent execution of hepatic functions. These interactions are subject to experimental validation.

**Figure 4.**
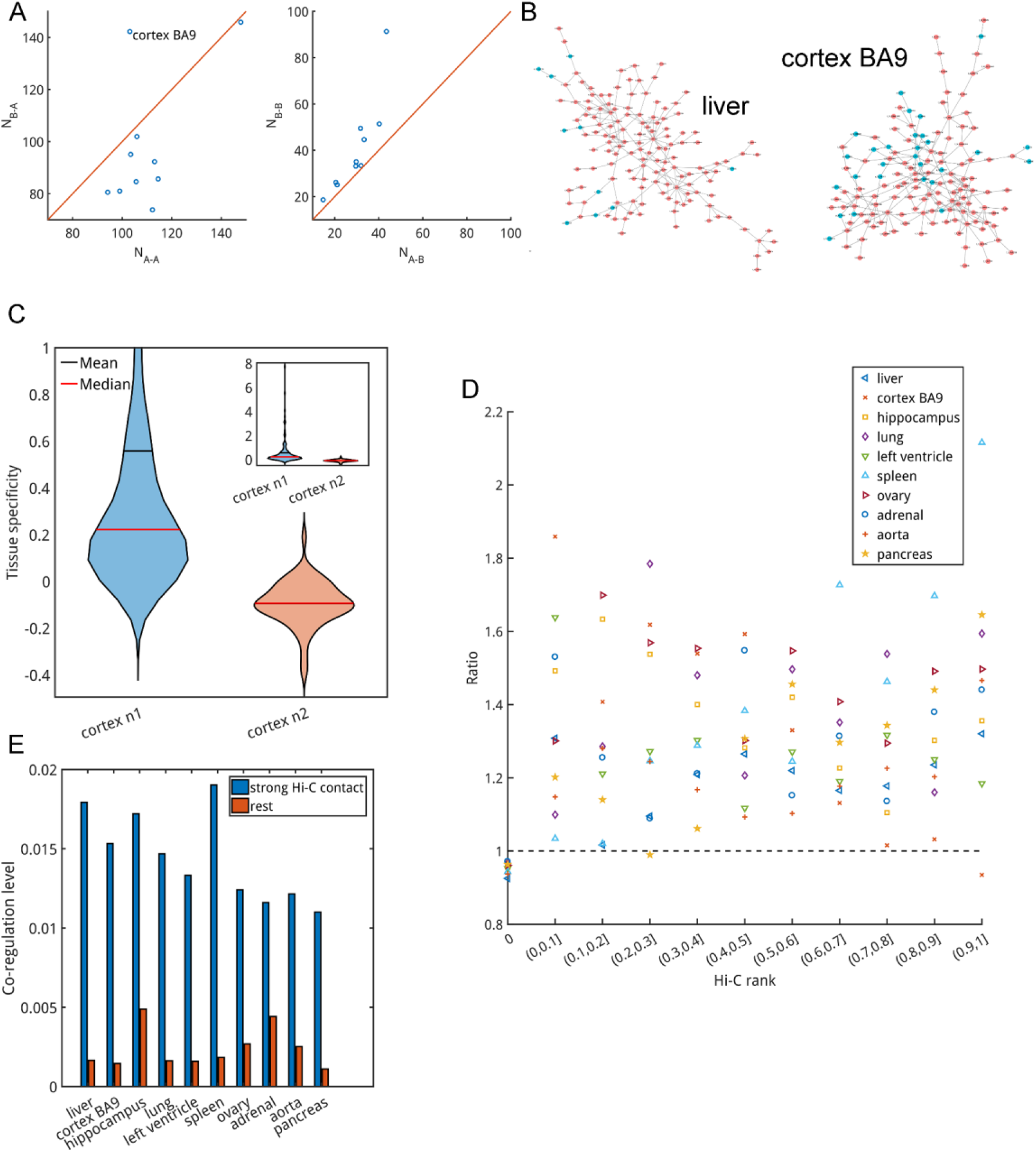
The interplay between gene pair expression coordination and chromatin structure. (**A**) Comparisons between N_A-A_ and N_B-A_ as well as N_A-B_ and N_B-B_ in the co-regulation networks of the ten cells. The only outlier is cortex BA9. (**B**) The two tissue-specific sub-networks for liver and cortex BA9 (Cytoscape software was used). (**C**) Comparison of the gene tissue specificity distribution between the two sub-networks in cortex BA9. n1, tissue-specific sub-network; n2, the generic sub-network. (**D**) The ratio between the Hi-C rank distributions of highly correlated gene pairs within chr1 and randomly chosen gene pairs 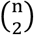, n is the number of genes of chr1). (**E**) The average correlation levels of highly contacted gene pairs compared to those that are not.

Next, we calculated the Hi-C rank between pairs of genes (see methods) to evaluate their relative Hi-C contact strength. The distribution of Hi-C rank of highly correlated gene pairs within a tissue at different Hi-C contact levels (11 level were used here) was calculated and compared with randomly chosen gene pairs from the gene list. The results given in **Figure 4D** show that in all tissues examined here, the ratio calculated between the Hi-C rank distributions of highly correlated gene pairs and randomly chosen pairs is always less than 1 for pairs of low Hi-C rank (equals 0) and greater than 1 for pairs of high Hi-C rank. Such a result shows that highly correlated gene pairs are more likely to form Hi-C contact than those of low expression correlations. Finally, we compared the gene expression correlation coefficients between gene pairs that are in high contact (see methods) to those that are not and found that the former has higher values (**Figure 4E and S25**). Therefore, gene pairs forming stronger spatial contacts are also more likely to be highly correlated in expression. These results all show a close association between chromatin structure and gene pair expression correlation, in addition to the expression levels of individual genes.

## Discussion and conclusion

In the present study, we try to connect the gene distribution in the linear and 3D genome to the establishment of tissue specificity. We integrated structural and genetic data to systematically study the relationship between gene expression and chromatin organization in somatic cells and observed a common role that 3D genome structure plays in gene regulation. Consistent with earlier studies (*30*), TSGs are found to distribute unevenly along the genome. In particular, CGI-deficient prairies significantly enrich genes of high tissue specificity compared with CGI forests, whereas the latter is abundant in non-tissue-specific genes and their overall tissue specificity is close to zero. More interestingly, the distribution of genes in prairie regions shows significant function-related mosaicity. Prairie but not forest domains tend to cluster genes of similar expression variations among different cell types (therefore probably similar biological functions). A small number of prairies were found to significantly enrich TSGs of similar functions and they are named functional modules. The assembly of genes of particular functions in linearly adjacent regions of low CpG density is likely to correlate with their co-activation in corresponding cells, possibly under the assist of specific TFs, and, equally important, co-repression in other tissues.

Besides the enrichment of TSGs, the relation between gene body CpG density and gene tissue specificity is also distinctly different between forest and prairie domains. For prairie but not forest genes, the tissue specificity is negatively correlated to gene body CpG density. In addition, a positive correlation was found between prairie gene body CpG density and compartment index, consistent with earlier studies (*24*, *25*) which showed a strong negative correlation between the repressive mark H3K9me3 and CpG density (or GC content). Therefore, the TSGs in prairies are prone to reside in heterochromatin and be repressed except in cells that they are specifically activated. Interestingly, many highly transcribed prairie TSGs remain in compartment B regions but are intermingled to a larger extent with forest regions in cells where they are highly expressed than in other cell types. Therefore, prairie genes appear to be activated through a tissue (cell)-specific interaction with the forest domains and a movement towards compartment A. The functional modules as a whole also display the strongest domain-mixing (between forests and prairies) in their corresponding tissues. Such a trend is also observed for the activation of forest TSGs, but to a lesser extent. Accordingly, we found that the activation of forest TSGs relies less on the intermingling with forest regions or moving to compartment A compared with prairies, indicating a more complex mechanism. We also found that prairie genes of high tissue specificities or expression levels tend to spatially contact with genes of similar tissue specificity or expression level in forests, which was reflected by the positive correlation between the tissue specificity (expression level) of prairie genes and the mean tissue specificity (expression level) of its highly contacted forest genes.

TFs have been widely reported to dynamically mediate chromatin structure. For example, TF binding sites can form clusters over long genomic distances (*21*, *22*). CTCF was proposed to participate in the TAD formation through a loop extrusion model (*34*, *35*) and was involved in tethering genome regions to nucleolus (*36*), indicating its capacity in chromatin repositioning. YY1, similar to CTCF, can form dimers to enhance interactions between enhancers and promoters (*37*). C/EBPα and OSKM were suggested to drive A-B compartment switch and TAD border insulation change during B-cell reprogramming (*13*). We thus speculate that TFs may also play a non-negligible role in mediating cell specific F-P interactions. In fact, our analyses did identify many F-P gene pairs, characterized by strong spatial contacts and high expression correlation levels (possibly sharing TFs (*38*)) for each tissue studied. Furthermore, the expression levels of a number of TFs tend to correlate with not only the expression level of these individual forest and prairie genes but also their correlation coefficient (**Table S3**), strongly suggesting a co-regulation of the forest and prairies genes by these TFs. For example, in liver, *CYP4A11* (forest gene) and *CYP2J2* (prairie gene) are highly correlated and in strong spatial contact and one liver-specific TF, NR1I3, correlates in expression with both *CYP* genes in liver and their correlation coefficients of expression. Together, these results are consistent with TFs interacting with both forest and prairie genes and especially facilitating the movement of the latter to a more transcriptionally active (forest) environment for regulation. This mechanism is consistent with the phase separation model in which the multivalent interactions of specific proteins can induce their condensation and subsequently form liquid-like droplets, resulting in the co-localization of genes. We expect that more studies focusing on the ChIP-seq data of such specific TFs will help test this prediction. In fact, a recent *in vitro* and *in silico* study has revealed that the DNA sequence features, including TF target site valence, density and affinity, play a vital role for driving condensation and both TF-DNA and TF-Mediator interactions determine the formation of condensates (*39*). Besides TFs, other molecules such as noncoding RNA may also play important roles in the spatial co-localization of genes.

Current analyses of the gene co-regulation network showed that in general genes are more likely to correlate in expression with other genes of the same than with genes of different compartment types. We then constructed a new type of gene networks, the edges of which represent not only high co-regulation level but strong spatial proximity, to investigate the spatial interactions of co-regulated gene pairs. We found that the interactions between forest and prairie genes in the cell-specific sub-network in liver and cortex BA9 do reflect the overall chromatin organization features, indicating a possible association between gene co-regulation and 3D structure. Compared to the standard gene network constructed by considering only gene expression data (*17*, *40*), this new network also contains information on the spatial interaction patterns of co-regulated genes and may be used in identifying tissue-specific gene clusters in the nuclear. Furthermore, we found that the co-regulated gene pairs are more likely to form Hi-C contact and the expression correlation coefficients of gene pairs in strong spatial contact are on average higher than others. These results illustrate the important roles chromatin organization plays not only in gene expression but also in gene regulatory network formation.

One intriguing discovery from the analysis of the gene network for cortex BA9 sample is that genes associated with neuro and metabolism functions are tightly coupled (co-regulated) and co-localized with each other. Namely, metabolic-related genes not only co-regulate in expression with but are also physically close to neuro-activity related genes. For example, *FABP3*, playing a role in the metabolism of long-chain fatty acids and lipoprotein, directly connects two other genes, *HPCA* and *SH2D5*. *HPCA* encodes a neuron-specific calcium-binding protein, which is thought to contribute to the function of neurons in the central nervous system (CNS), and the protein encoded by *SH2D5* is known to participate in the regulation of synaptic plasticity. *ST3GAL3*, which is associated with the metabolism of glycosaminoglycan and glycosphingolipid, connects two genes *EPHA10* and *SNAP47*. The protein encoded by *EPHA10* acts as a mediator of axon guidance and cell-cell communication in neuronal and epithelial cells. *SNAP47* is related to syntaxin binding. Notably, it is known that the pathway of glucose metabolism is highly relevant to neurodegenerative disorders such as Alzheimer disease (*41*). As another example, *SLC45A1*, which encodes glucose transporter in the brain, is correlated to and in strong spatial contact with *HTR6* that encodes a G protein-coupled protein, which regulates cholinergic neuronal transmission in brain and spatial memory. *SLC44A5*, similar to *SLC45A1*, is connected to *RGS7* which regulates G protein signaling and synaptic vesicle. *PRKCZ*, related to glucose transport upon insulin treatment, connects *GABRD* that mediates neuronal inhibition. These and other examples show that the execution of normal brain functions heavily depends on these cross-function (and cross forests and prairies) interactions (both in space and in expression regulation). Interestingly, previous studies showed that a possible connection does exist between neurodegenerative diseases, e.g., Alzheimer disease, and the digestive system, such as the metabolism of lipid (*42*, *43*) and bile acids (*44*).

In our previous study, we found compartments A and B are mainly consisted of forests and prairies, respectively (*23*), indicating the strong DNA sequence-dependence in chromatin compartmentalization. In contrast to the composition of compartments A and B which can significantly vary in different cells (*26*), the forest and prairie, identified solely based on sequence property, are nearly constant along the different cell stages, such as early embryo development, cell differentiation and senescence. In this study, we noticed the extent of F-P intermingling is strongly concordant with the activation or repression of genes related to the establishment of tissue specificity, providing a simple and uniform framework (the forest-prairie domain segregation) for understanding the achievement of cell identity and the evolution of chromatin structure among different biological stages. Besides individual genes, our results also provide insights into the interplay between gene pair expression coordination and genome organization.

In summary, our study showed that, consistent with earlier observations (*30*), the TSGs related to one cell type and genes of similar functions tend to cluster along the human genome, especially in CGI-poor and normally gene-poor prairie domains. The tissue specificity of TSGs in prairie, and to a much less extent, that in forest, anti-correlates to the corresponding gene body CpG density. In 3D chromatin structure, theses prairie TSGs relevant to one certain tissue need to interact spatially with gene rich forest domains in a tissue-specific manner for activation. The forest genes in tissue-specific spatial contact with these prairie TSGs also tend to possess high tissue specificity. In other tissues, the irrelevant prairies TSGs tend to move to a more compartment B, and thus presumably more repressive, spatial environment. Accordingly, in the absence of tissue-specific forest-prairie contacts, the TSGs also tend to have low expression levels, providing a mechanism of establishment and maintenance of tissue specificity given the complex but common DNA sequence shared by all tissues.

## Supplementary Information

Is available in the online version of the paper.

## Acknowledgement

This work was supported by National Natural Science Foundation of China (21821004, 21873007) and National Key R&D Program of China (2017YFA0204702).

## Author contributions

Yi Qin Gao designed the research. Hao Tian performed the research with the help of Ying Yang, Sirui Liu and Hui Quan. Hao Tian and Yi Qin Gao wrote the manuscript.

