## Supplementary Information for "Toward an understanding of the relation between gene regulation and 3D genome organization"

### Supplementary Text

**Somatic cells analyzed in this study**

Ten somatic cells, liver, cortex BA9, hippocampus, lung, left ventricle, spleen, ovary, adrenal, aorta and pancreas, were mainly used in this study to investigate the relation between gene regulation and genome organization, as they were performed both Hi-C (*1*) and RNA-seq (*2*) experiment. Notably, Sonawane *et al*. provided normalized RNA-seq data of nearly 40 tissues (*2*), which was further utilized in calculating gene tissue specificity and identifying TSGs in certain cell type.

**Hi-C data alignment among different cells**

Due to the different Hi-C data sizes of different cells, we discarded the Hi-C elements that were smaller than $\alpha^t$, the value of which is given in **Table S1**, aligning the Hi-C data of different cell types to a similar scale so that they can be compared.

### Supplementary Tables

**Table S1. Threshold and size of Hi-C data in ten cells**

| | Liver | Cortex | Hippocampus | Lung | LV | Spleen | Ovary | Adrenal | Aorta | Pancreas |
|---|---|---|---|---|---|---|---|---|---|---|
| $\alpha^t$ | 1.8960 | 0.4272 | 0.4790 | 0.2249 | 2.0133 | 0.4049 | 0.3242 | 0.3620 | 1.3111 | 0.3754 |
| Before | 4371504 | 1588909 | 1780322 | 1040669 | 2626072 | 1533076 | 1406427 | 1385049 | 3549393 | 1476175 |
| After | 1187231 | 1563662 | 1726100 | 1034985 | 1488972 | 1507728 | 1393523 | 1367474 | 1569774 | 1432612 |

*"Before" means the number of non-zero elements in the Hi-C data of each cell and "After" represents the number of elements, the value of which is larger than $\alpha^t$. LV=left ventricle.

**Table S2. Functional module location**

| Cell name | Chromatin | Location/bp |
|---|---|---|
| Liver | Chr4 | 69215613-72897763 |
| Spleen | Chr19 | 54711241-55450974 |
| Adrenal | Chr6 | 52535852-52859665 |
| Pancreas | Chr1 | 101702596-108507545 |

**Table S3. Examples of high order correlation between F-P gene pairs and TFs in liver**

| **Forest gene** | **Prairie gene** | **TF** |
|---|---|---|
| *CYP4A11* | *CYP2J2* | NR1I3 |
| *ACADM* | *GBP7* | NR1I3 |
| *AK4* | *GBP7* | NR1I3 |
| *TMEM56* | *DPYD* | ESR1 |
| *CA14* | *FMO3* | ARID3C |

*In each example, the forest and prairie genes are not only highly correlated but forming strong spatial contact in liver. Furthermore, the expression level of corresponding TF correlates with not only the individual forest and prairie genes in liver, but the correlation level between them.

#### Supplementary Figures

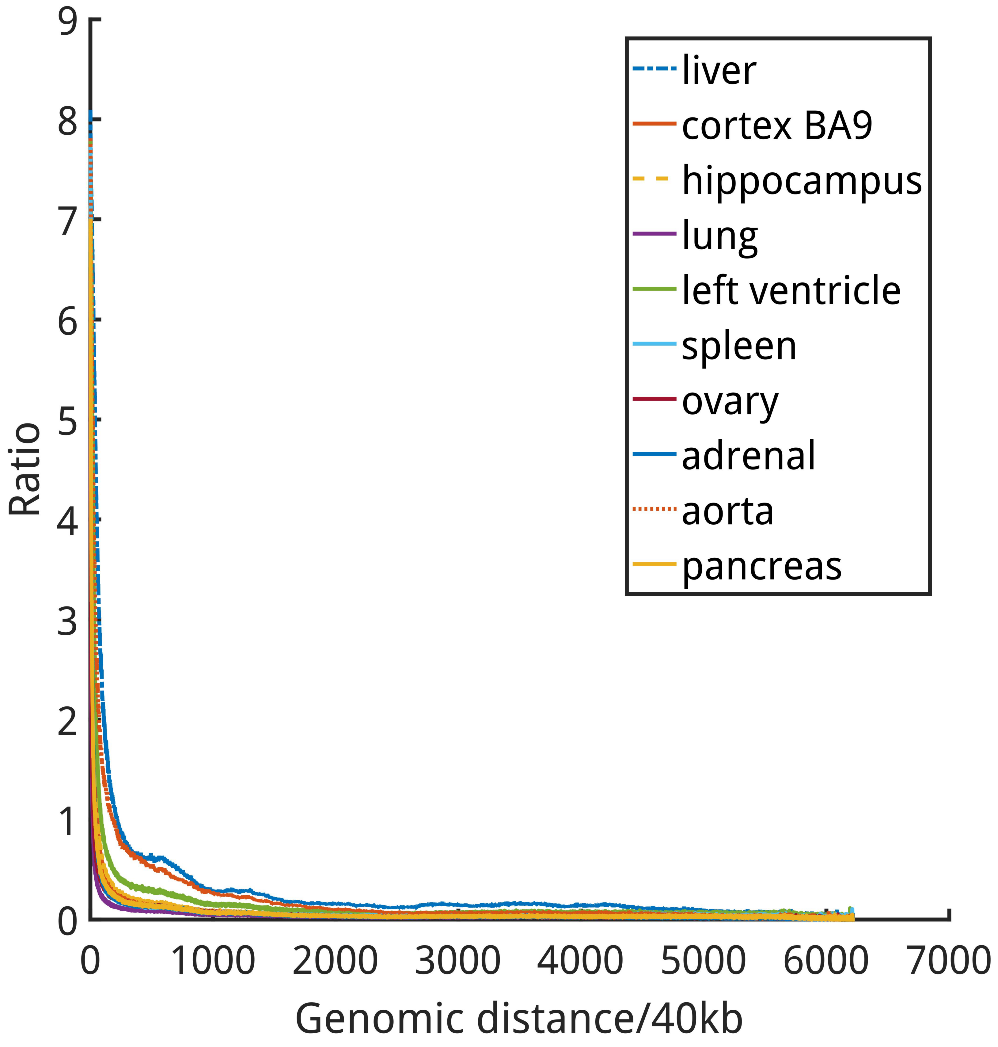


**Figure S1. The ratio calculated between the number of non-zero and zero Hi-C elements given one certain genomic distance.**

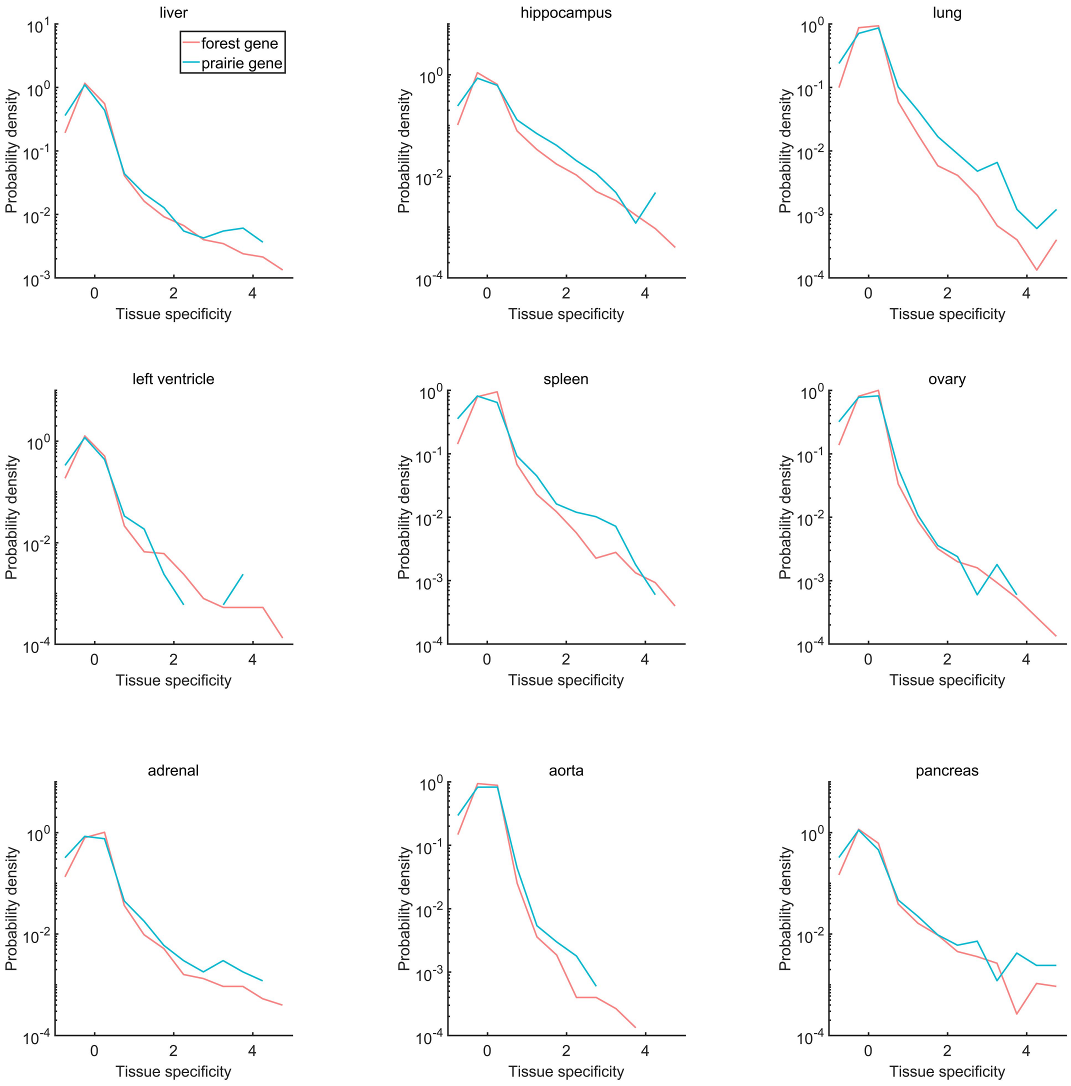


**Figure S2. The probability density distribution of tissue specificities of forest and prairie genes in nine cells.** The range of gene tissue specificity was chosen as [-1,5], resulting in the missing of several genes in some cells (the total number of forest and prairie genes is 18413).

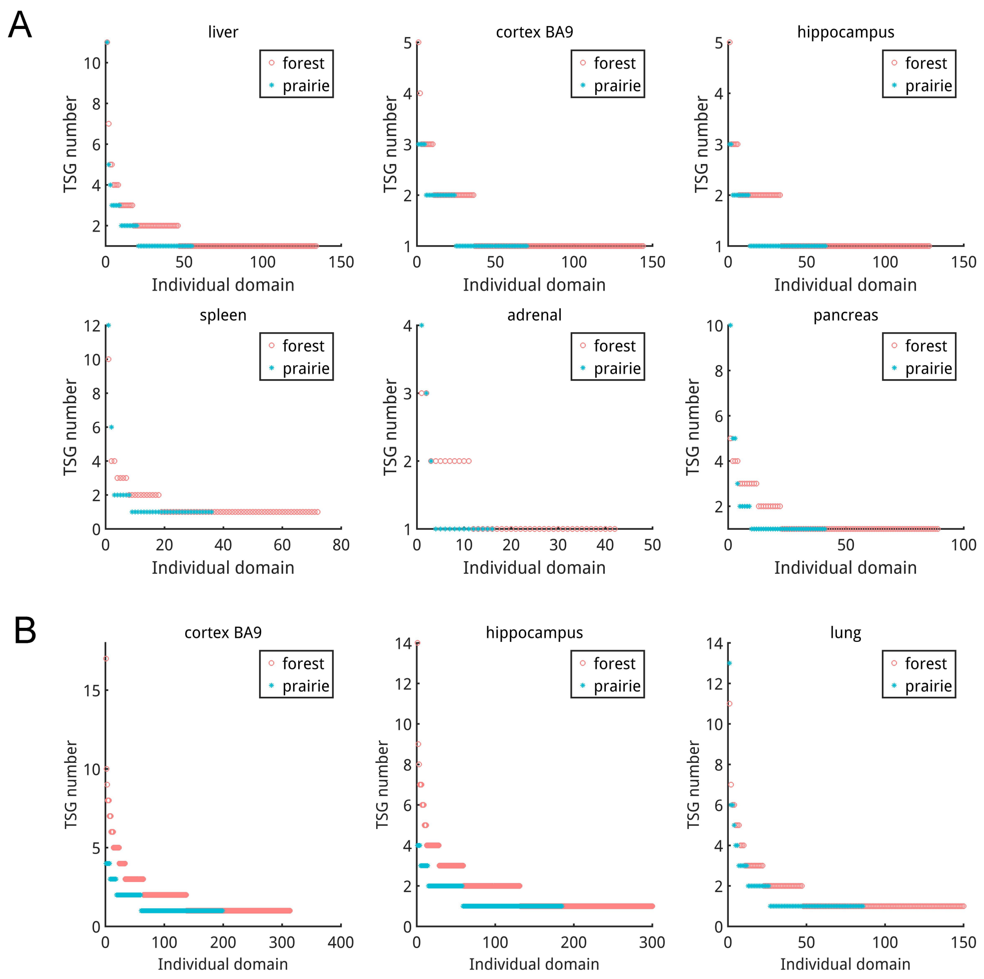


**Figure S3. The number of TSGs relevant to one tissue within individual forest and prairie domains.** We identified several prairie domains, named functional modules that significantly harbor TSGs belonging to one tissue, in liver, spleen, adrenal and pancreas. In contrast, two brain tissues, cortex BA9 and hippocampus, exhibit the broader and more uniform distribution of related prairie TSGs and thereby do not possess functional modules under the two criteria in TSG identification: $s_i^t > 2$ (**A**) and $s_i^t > 1$ (**B**).

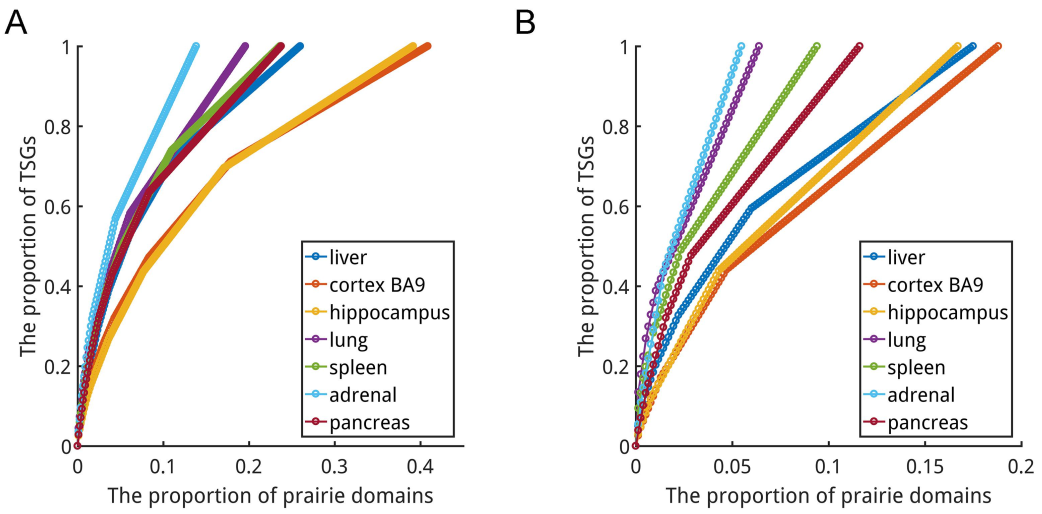


**Figure S4. "Saturation curve" for forest TSGs under the criteria $s_i^t > 1$ (A) and $s_i^t > 2$ (B), corresponding to Figure 1C.**

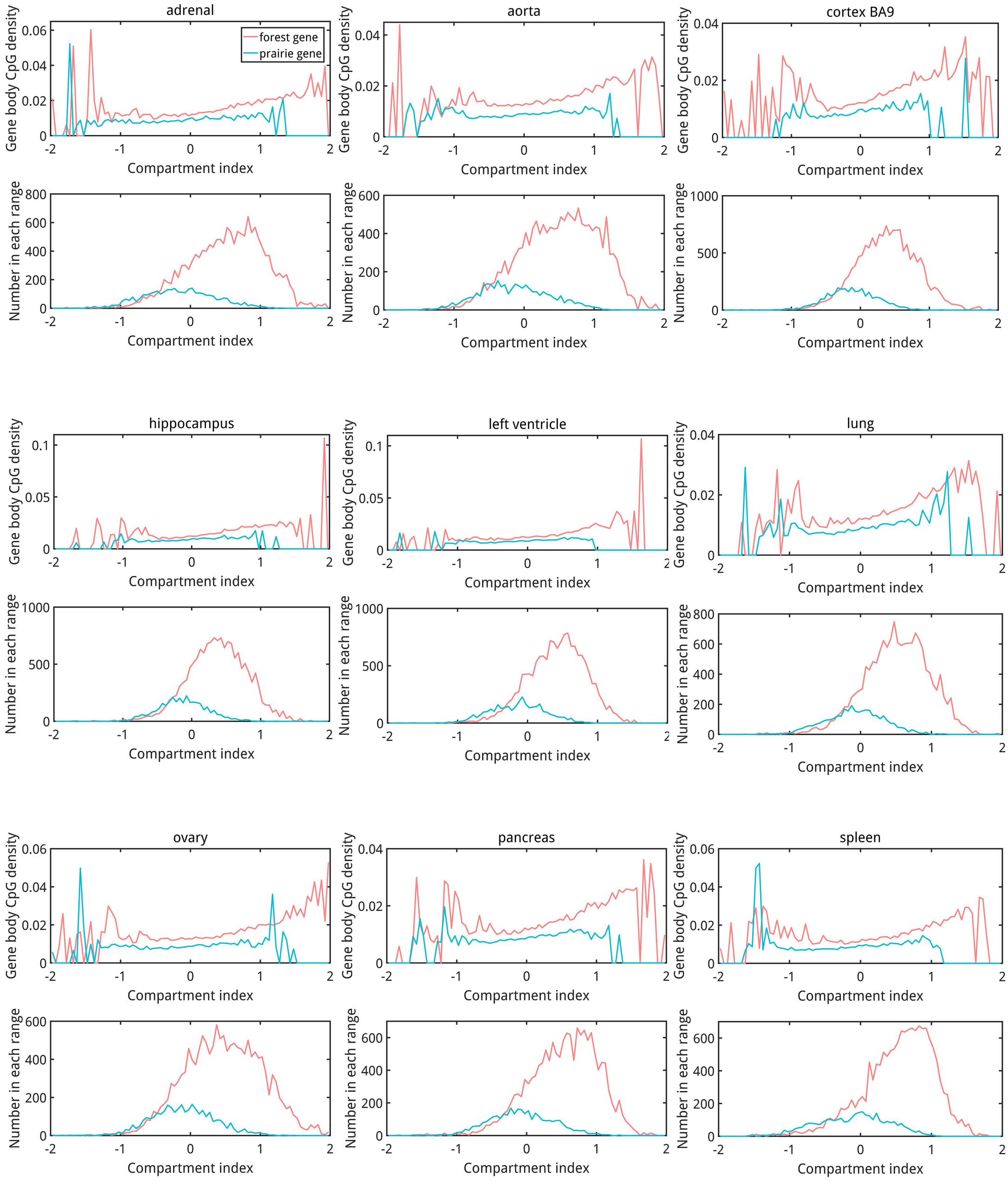


**Figure S5. The relation between compartment index and gene body CpG density in nine cells, corresponding to Figure 2B.** The range of compartment index was chosen as [-2, 2], resulting in the missing of several data points in some cells.

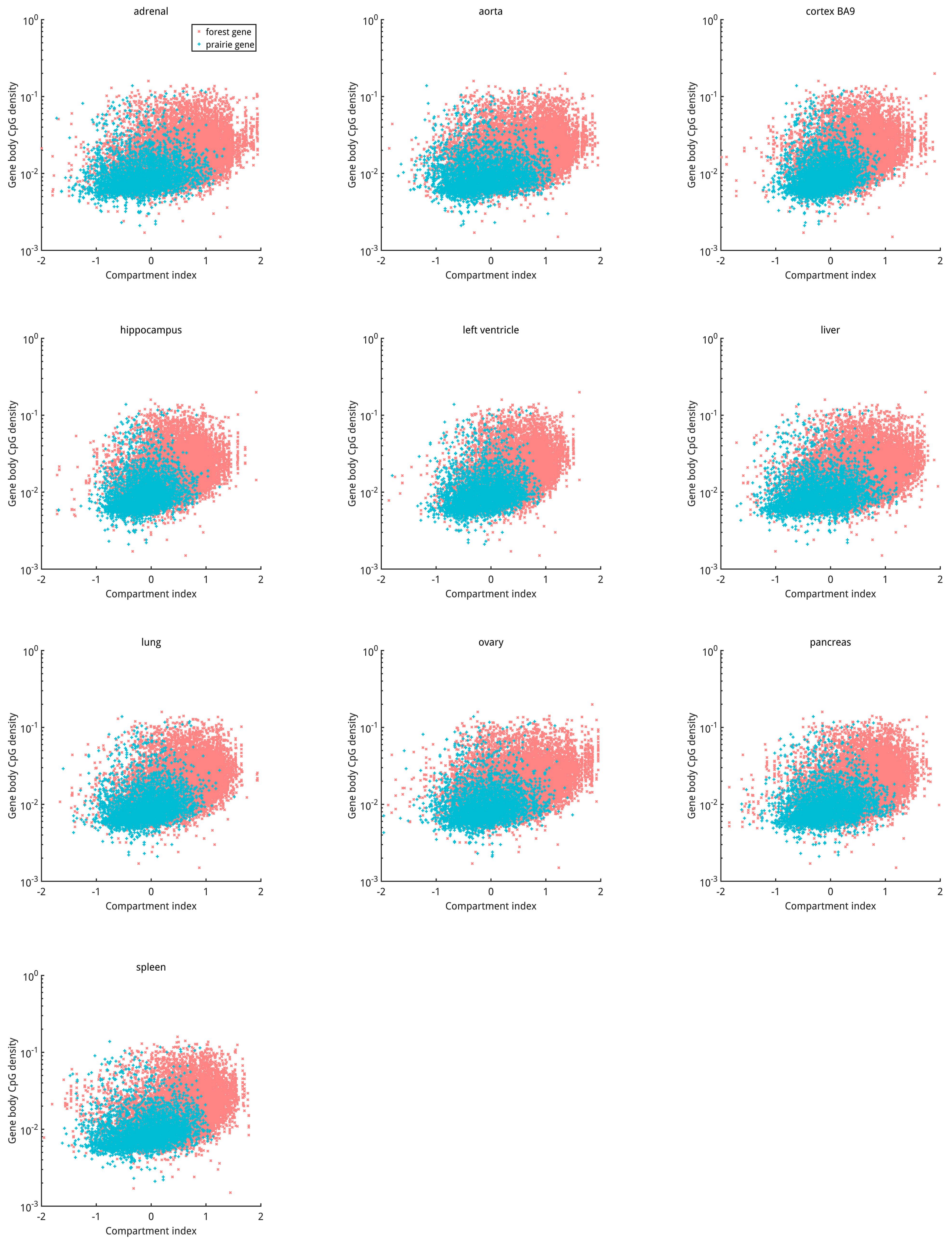


**Figure S6. Scatter plot for the relation between compartment index and gene body CpG density.**

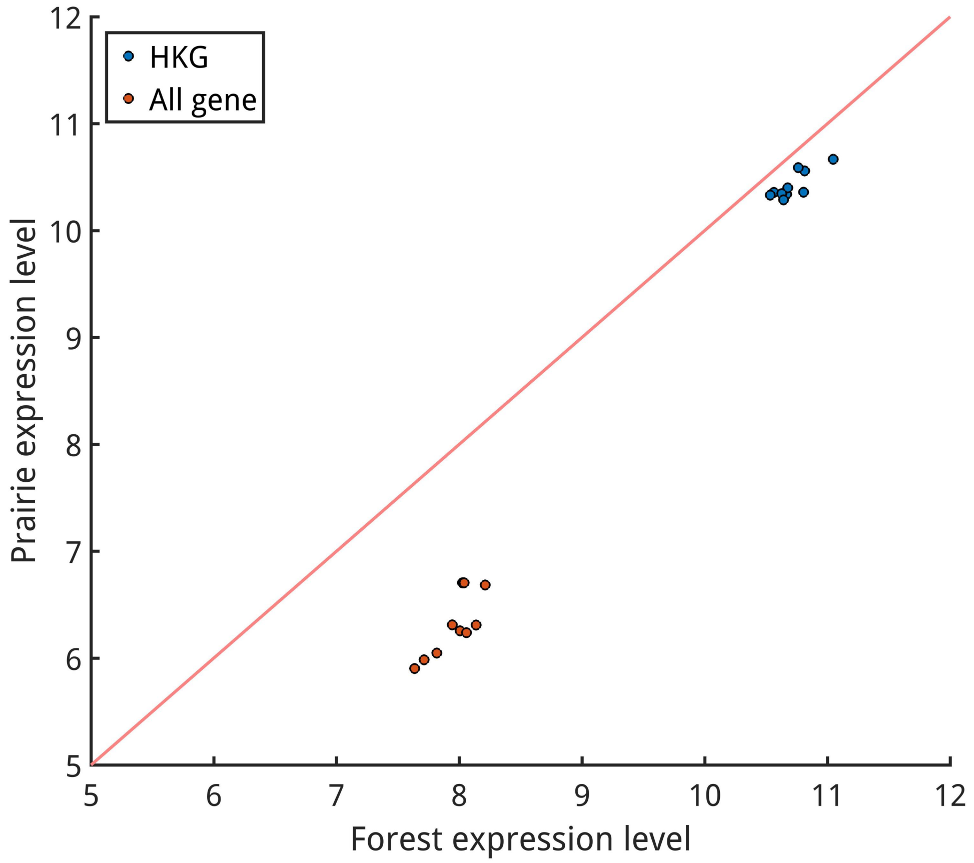


**Figure S7. The difference of gene/HKG expression levels between forest and prairie.** Ten cells, liver, cortex BA9, hippocampus, lung, left ventricle, spleen, ovary, adrenal, aorta and pancreas, were considered and each data point represents the average expression level of forest/prairie genes/HKGs in one of the ten cells.

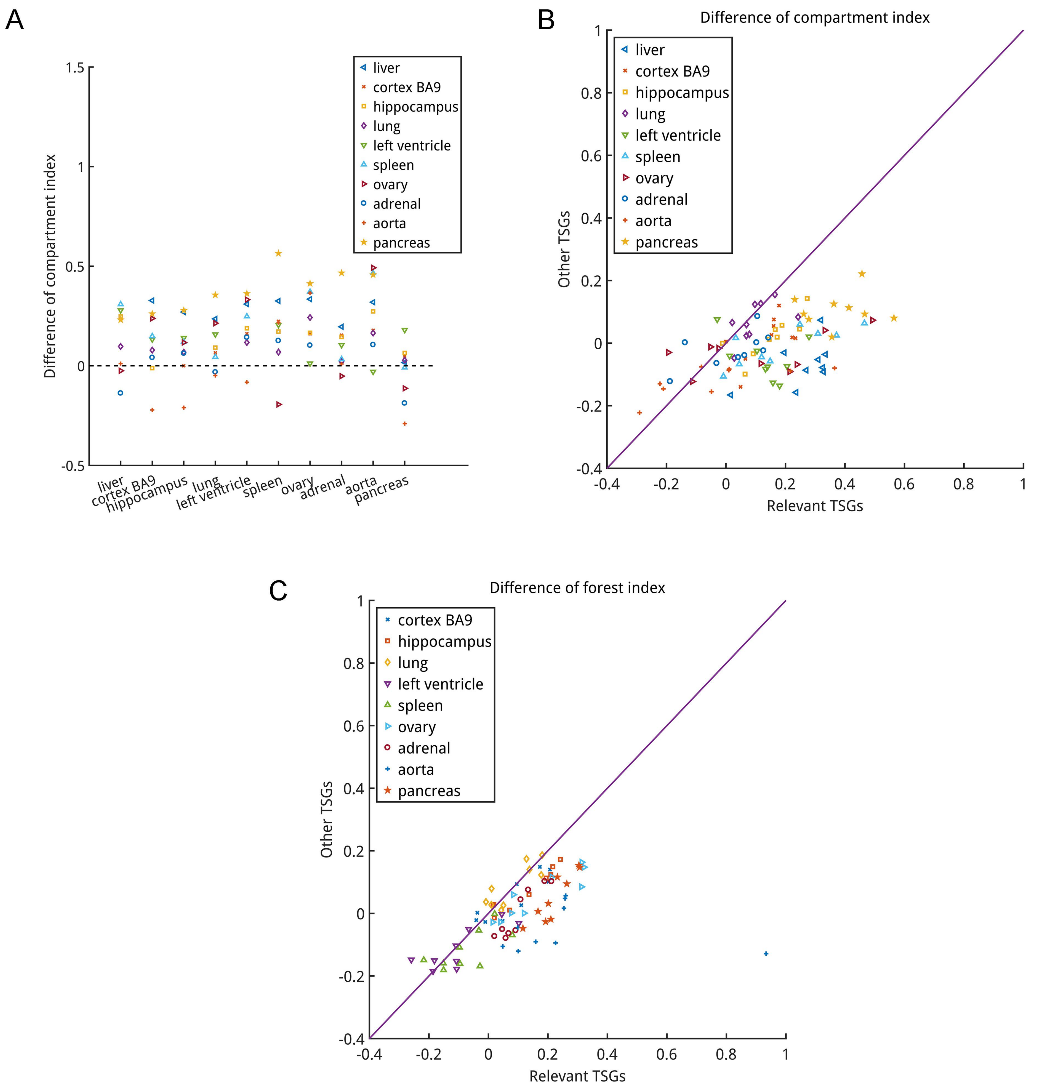


**Figure S8. The regulation of prairie TSGs is strongly associated with 3D genome organization. (A)** Compartment index change of one group of prairie TSGs belonging to one certain cell type (e.g., liver prairie TSGs) from the nine control cells (e.g. spleen) to relevant cell (e.g. liver). The value behind each symbol represents the median compartment index difference between related cell (illustrated in legend) and control cell (labeled in x-axis) for related cell's prairie TSGs. Similar to forest index (**Figure 3B**), the generally positive value of compartment index change indicates that prairie TSGs enter a more transcription-active environment when needing to be highly transcribed. **(B-C)** Comparison of (B) compartment and (C) forest index changes between relevant TSGs (e.g. liver prairie TSGs) and other TSGs (TSGs of the remaining cells) when cell type transfers from control cell (e.g. spleen) to relevant cell (e.g. liver). In each figure the median values were shown. In such a process, we observed a higher and more positive value of compartment/forest index change for relevant TSGs, indicating a more favorable transcription environment they entered, compared to TSGs of the other cells. Together, these results emphasize the close link between high order chromatin structure and gene expression.

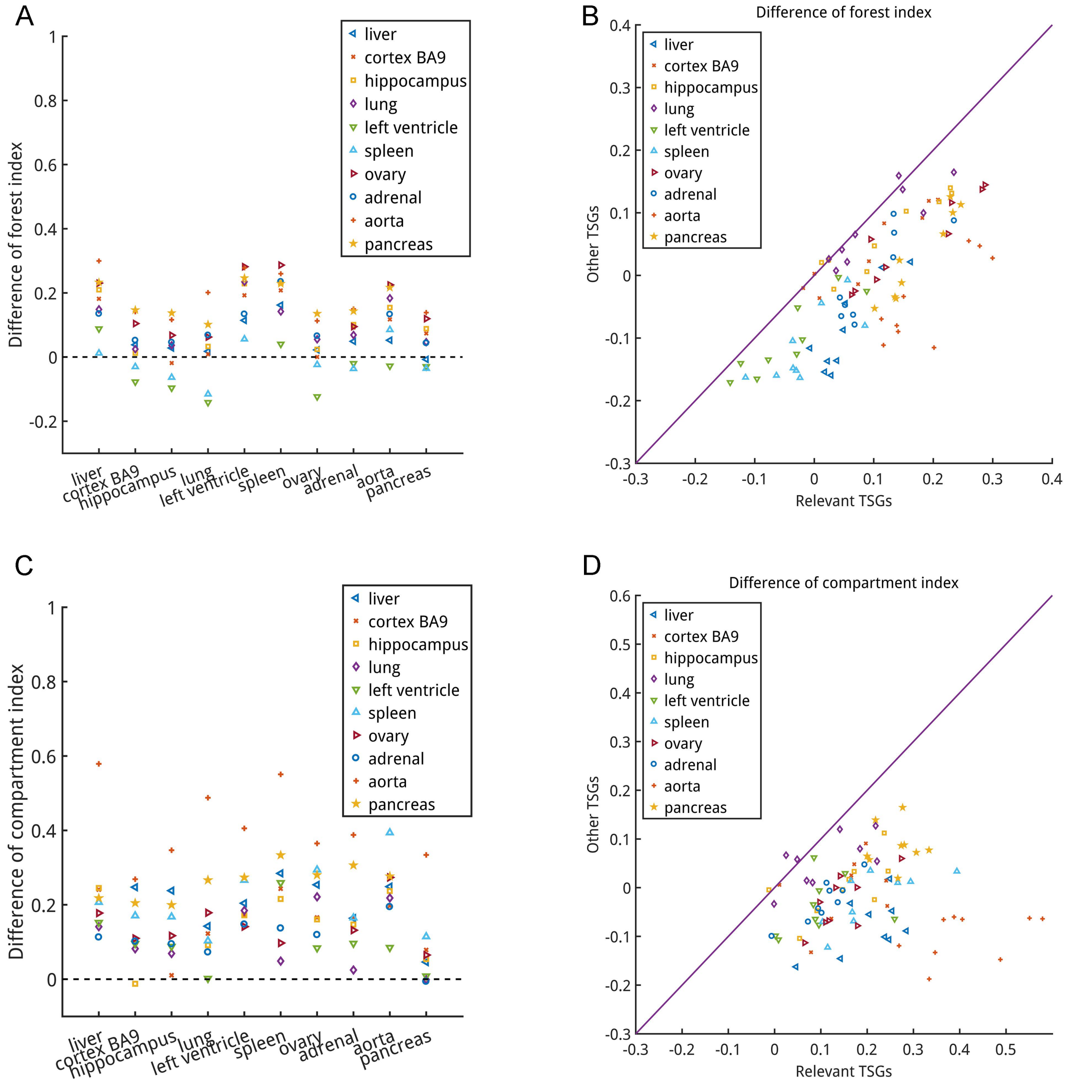


**Figure S9. A robustness test for the relation between prairie gene regulation and genome organization, corresponding to Figure 3B, 3C and S8.** The criterion we used here in TSG identification is $s_i^t > 1$.

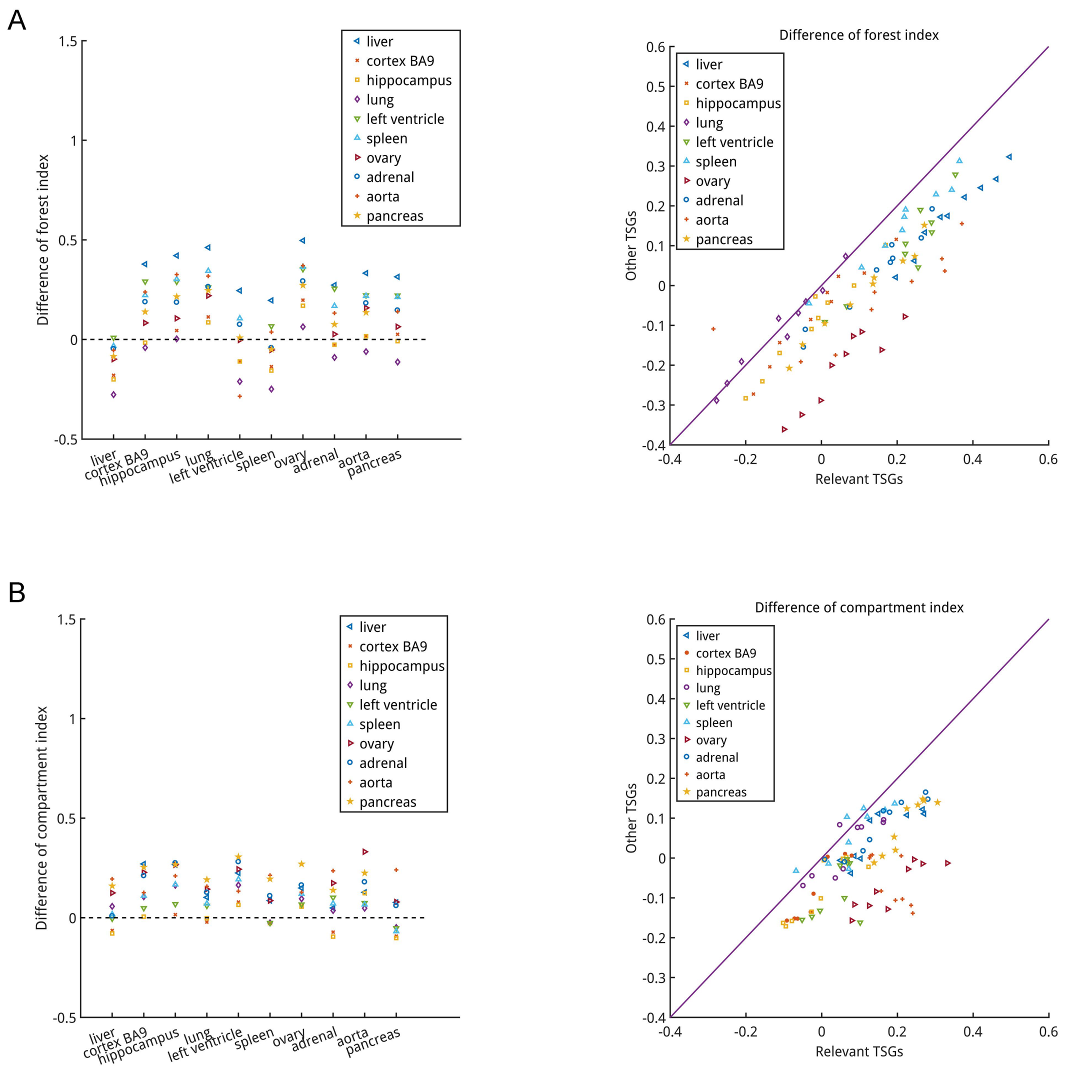


**Figure S10. The regulation of forest TSGs is also associated with genome organization. (A-B)** Akin to prairie TSGs (**Figure S8, S9, 3B and 3C**), the activation or repression of forest TSGs are also related to (A) forest and (B) compartment index changes, i.e., the chromatin structure reorganization (median values were shown).

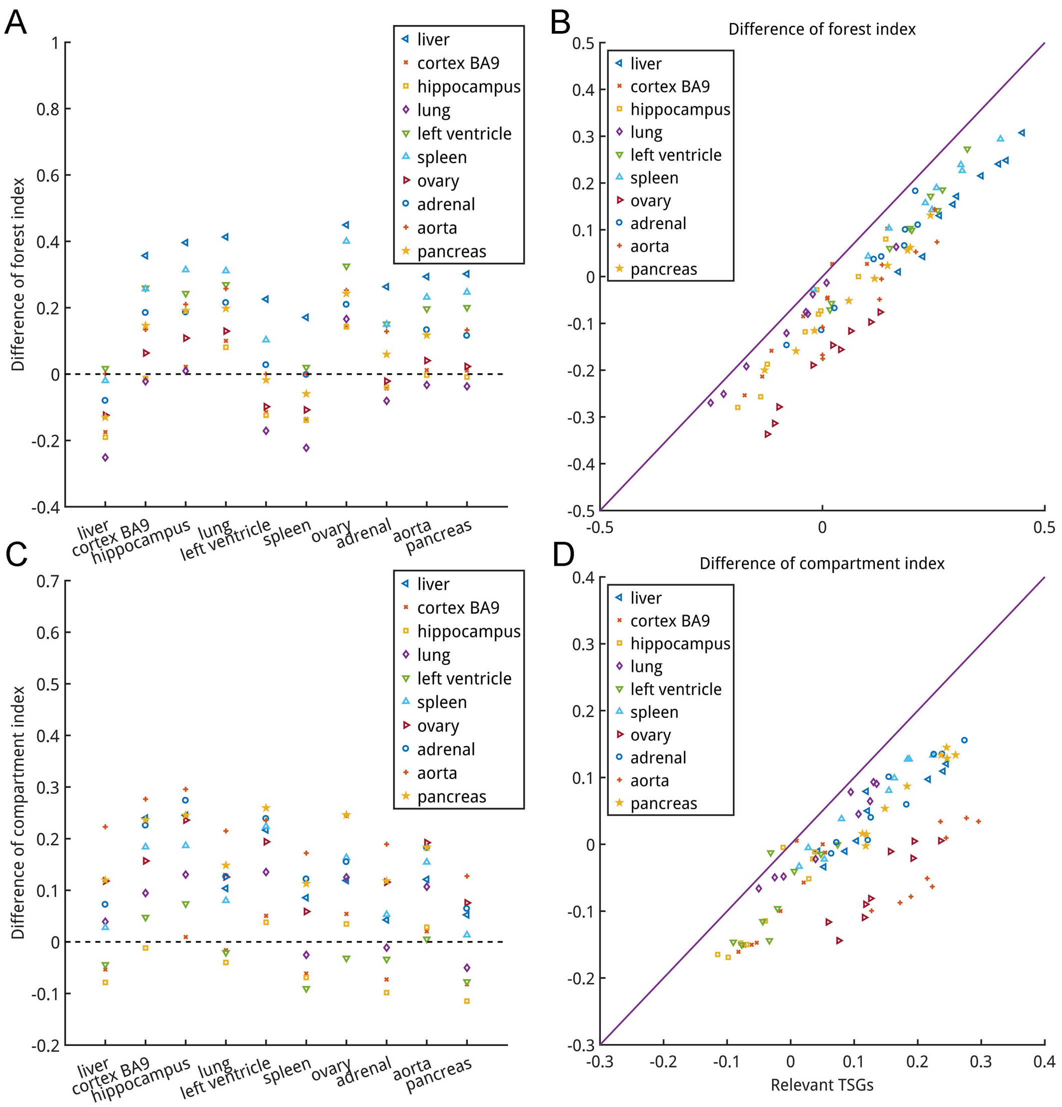


**Figure S11. A robustness test for the relation between forest gene regulation and genome organization, corresponding to Figure S10.** The criterion we used here in TSG identification is $s_i^t > 1$.

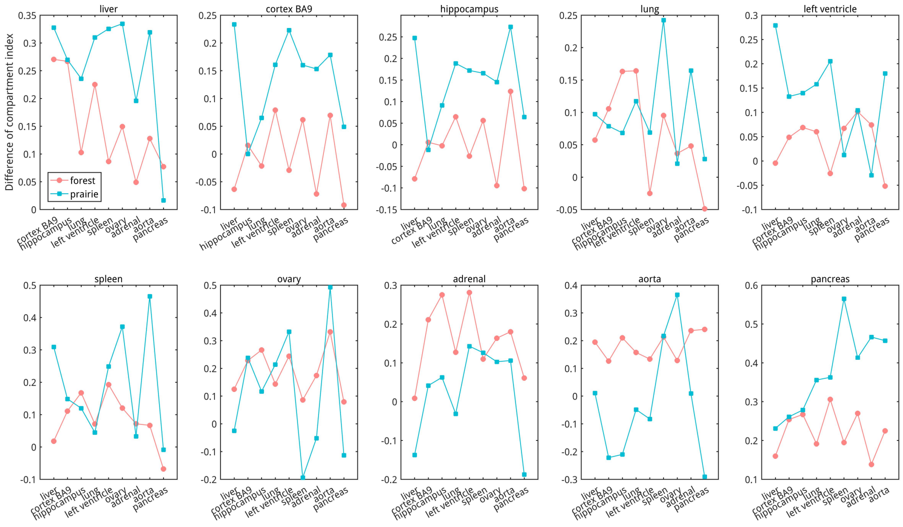


**Figure S12. The activation of prairie TSGs depends more on specific chromatin structure reorganization.** We further compared the compartment index changes of forest and prairie TSGs relevant to one cell type (e.g., liver) when cell type transfers from control cells (e.g., spleen) to relevant cell (e.g., liver) and found such extent of prairie TSGs is larger than forests in most cases. Each data point represents the median value of compartment index changes, corresponding to **Figure S8A** and **S10B**.

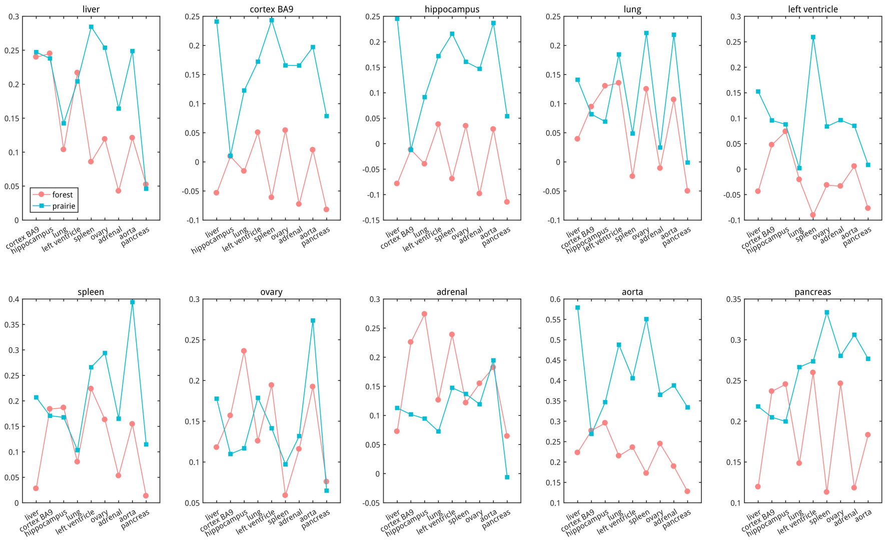


**Figure S13. A robustness test, corresponding to Figure S12.** The criterion we used here in TSG identification is $s_i^t > 1$.

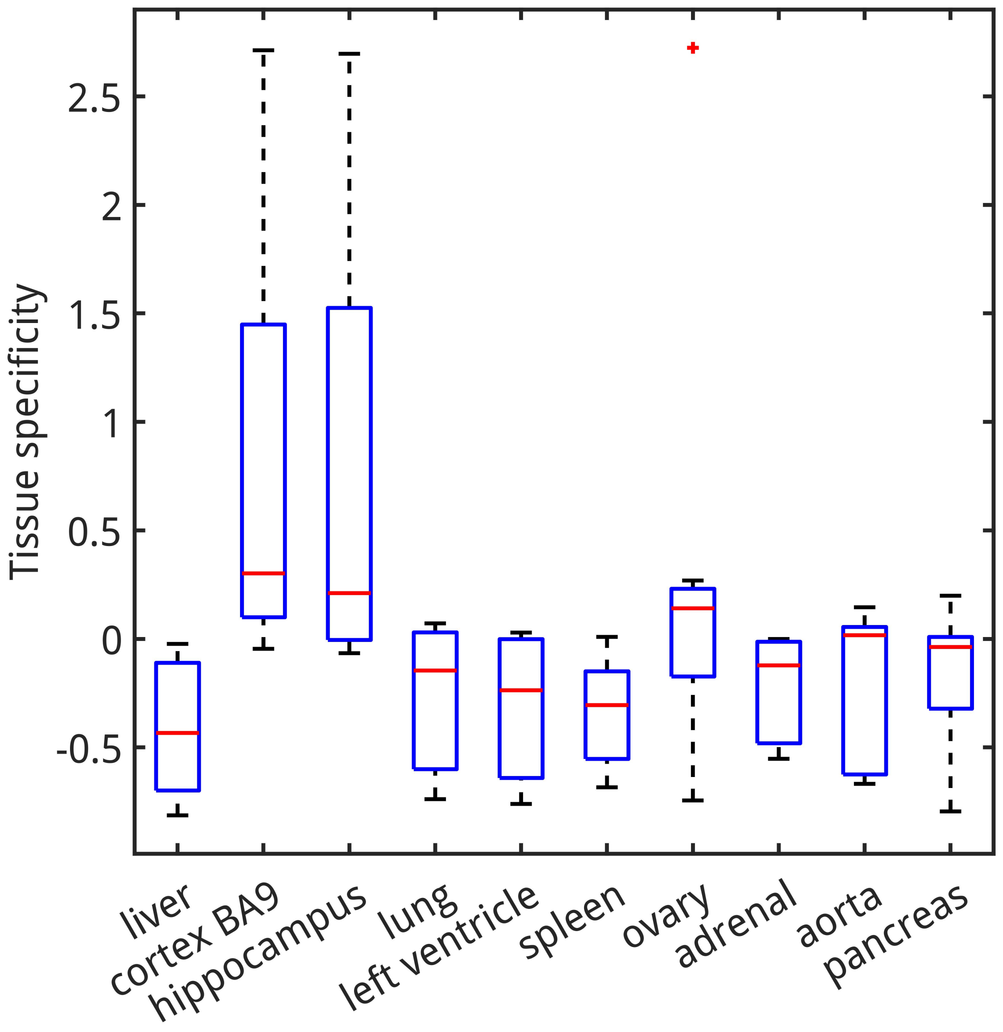


**Figure S14. The tissue specificities of eight representative forest genes showing the most negative correlation between gene expression level and forest index.**

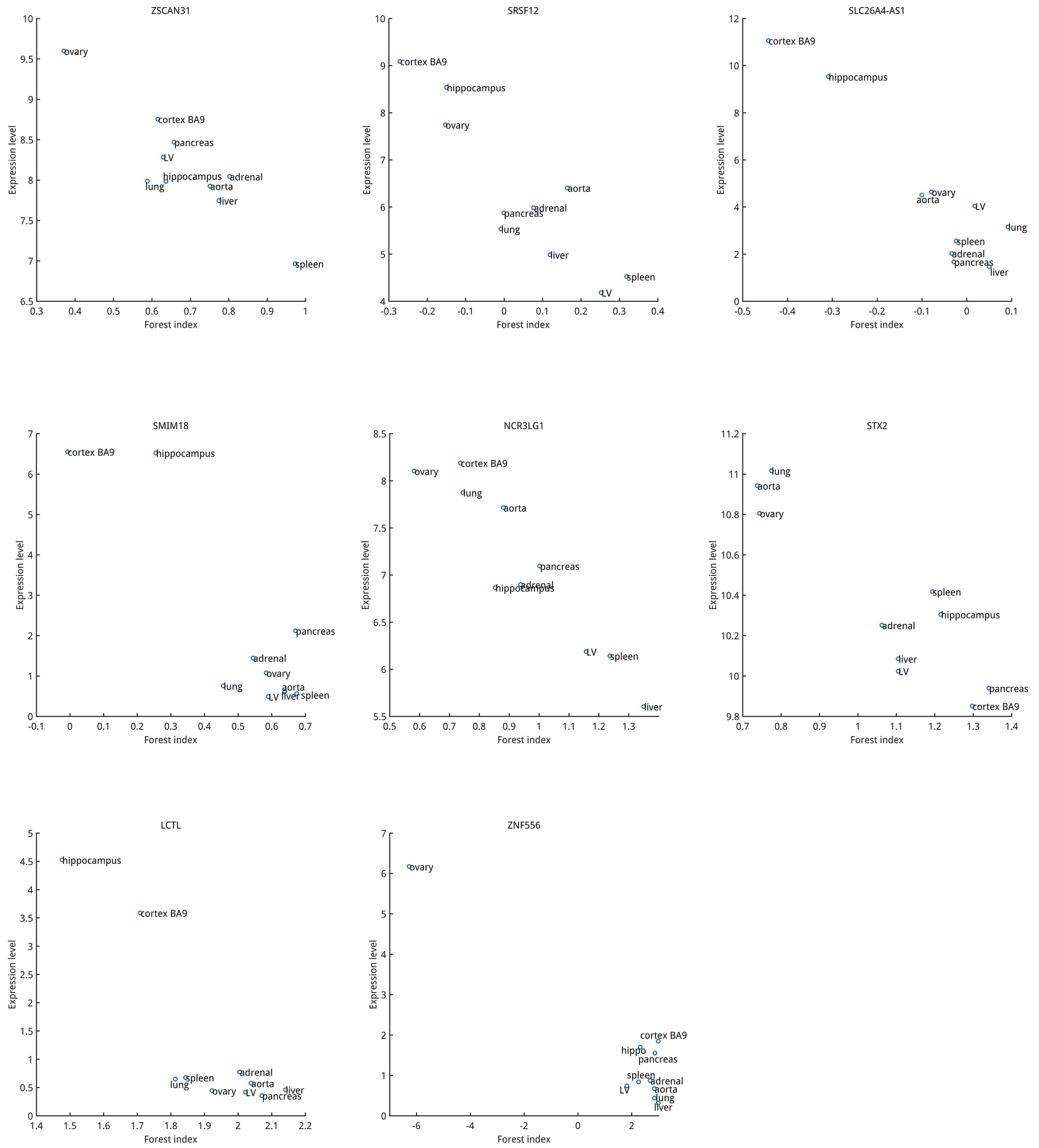


**Figure S15. Detailed information about the expression level and forest index of eight genes in ten cells, corresponding to Figure S14.** LV=left ventricle.

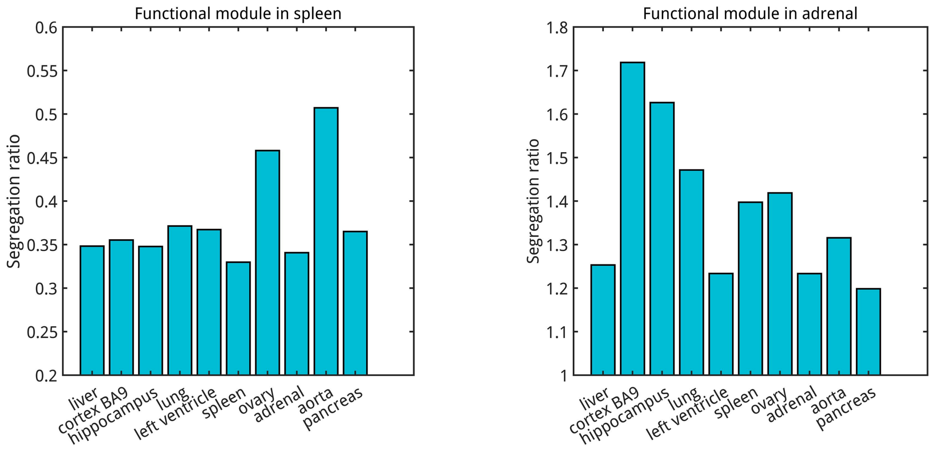


**Figure S16. The segregation ratio of spleen and adrenal functional modules in ten cells.** We noticed that the segregation ratio of adrenal functional module in adrenal is litter bigger than that in pancreas and then check the tissue specificity of genes residing in this domain in pancreas. The results revealed that three out of six genes also possess high tissue specificity in pancreas: *GSTA7P*, 8.73; *GSTA2*, 4.69; *GSTA1*, 1.80. Therefore, similar to adrenal, these genes also need to intermingle with forest regions for activation in pancreas.

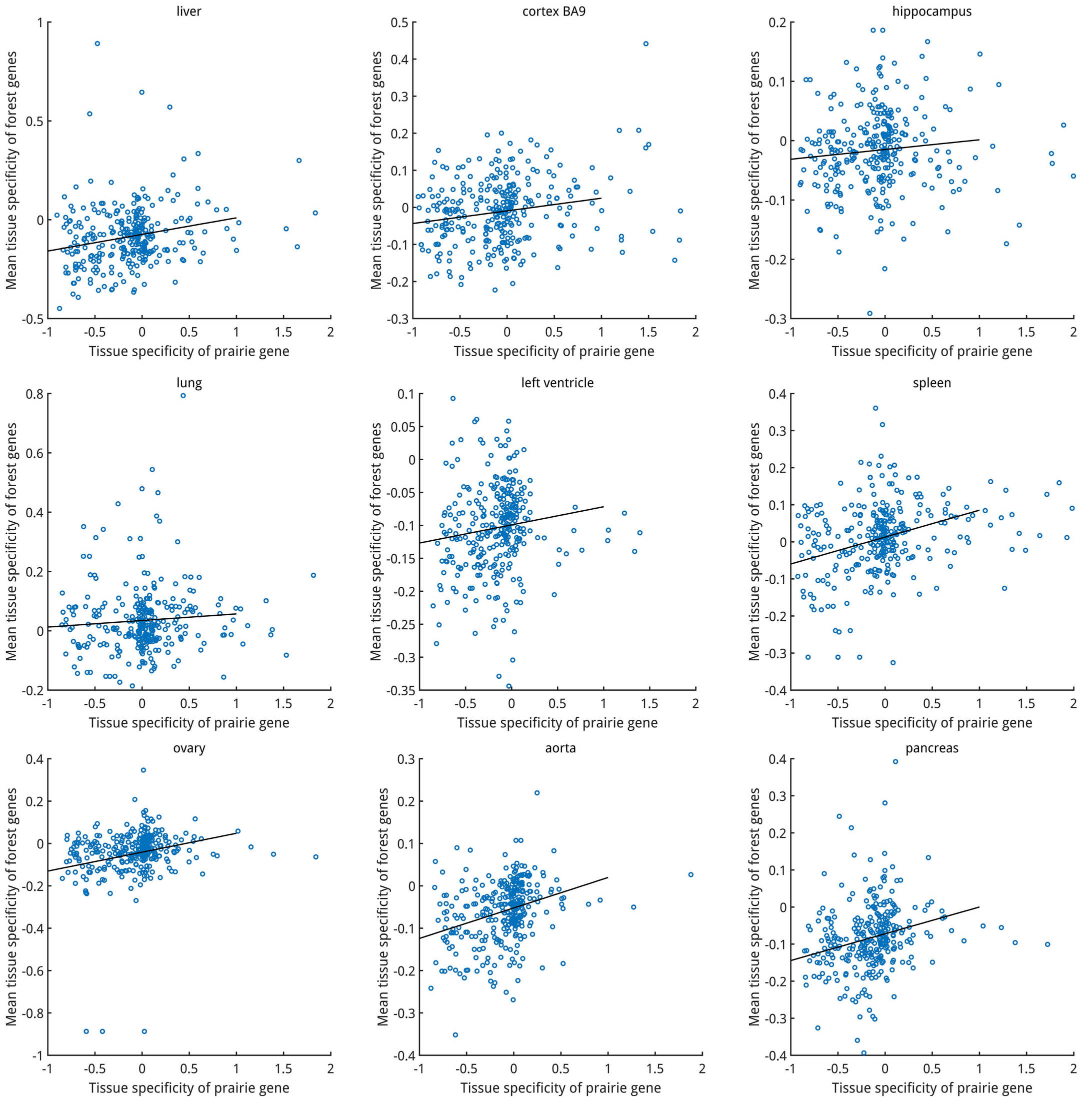


**Figure S17. The relation between tissue specificity of prairie gene and the mean tissue specificity of its highly contacted forest genes in nine cells.** All prairie genes of chr1 (the number is 317) were selected for calculation. When drawing these figures, the range of prairie gene tissue specificity (x-axis) was selected within $[-1,2]$ and 95.9%~100% prairie genes were retained in these nine cells.

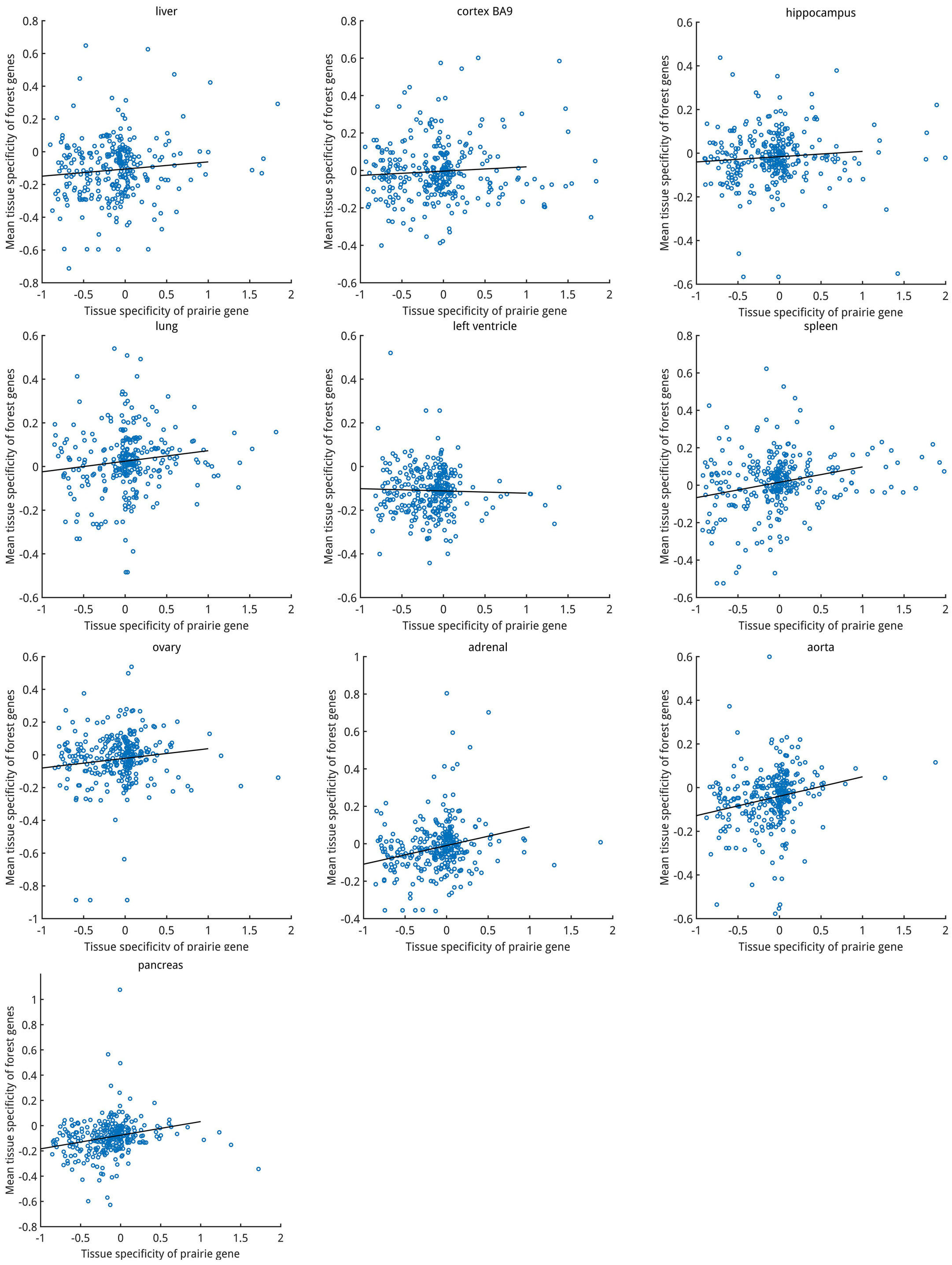


**Figure S18. A robustness test for the relation between tissue specificity of prairie gene and the mean tissue specificity of its highly contacted forest genes, corresponding to Figure S17.** Here the criterion we used in the identification of highly contacted gene pairs is 90$^{th}$ percentile (top10%, see methods).

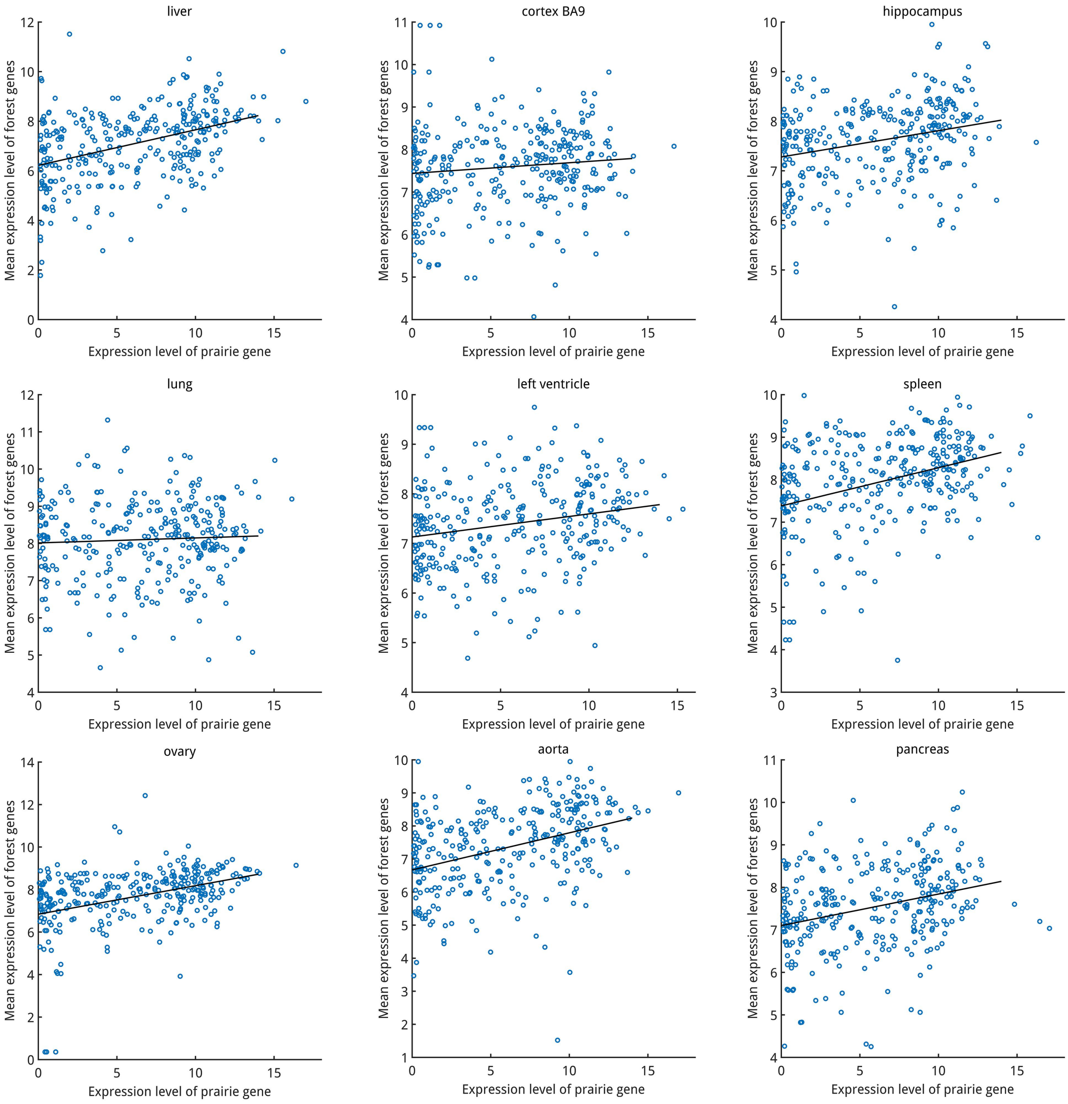


**Figure S19. The relation between expression level of prairie gene and the mean expression level of its highly contacted forest genes in nine cells.**

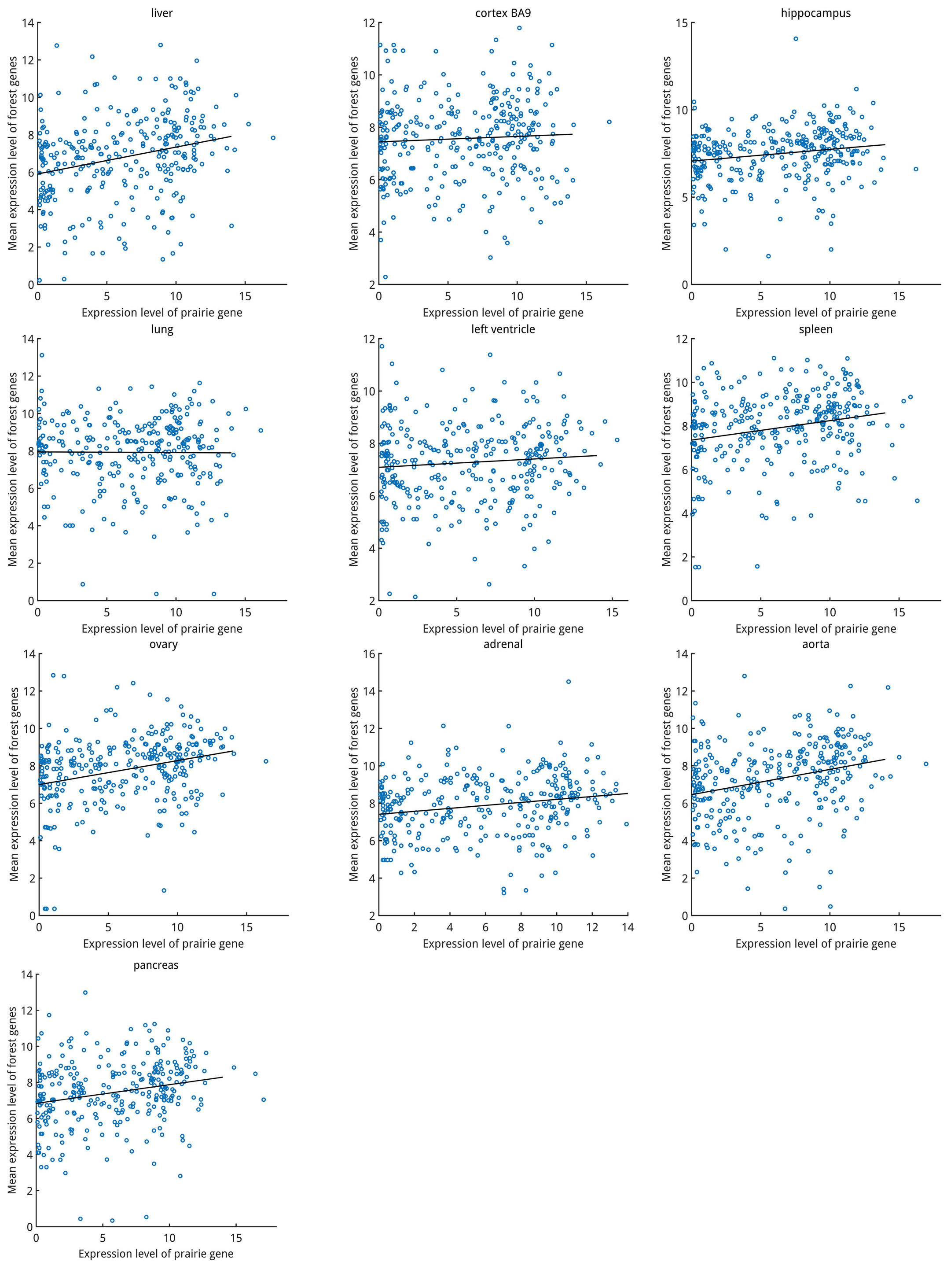


**Figure S20. A robustness test for the relation between the expression level of prairie gene and the mean expression level of its highly contacted forest genes, corresponding to Figure S19.** The criterion we used here in identifying highly contacted gene pairs is 90$^{th}$ percentile (top10%, see methods).

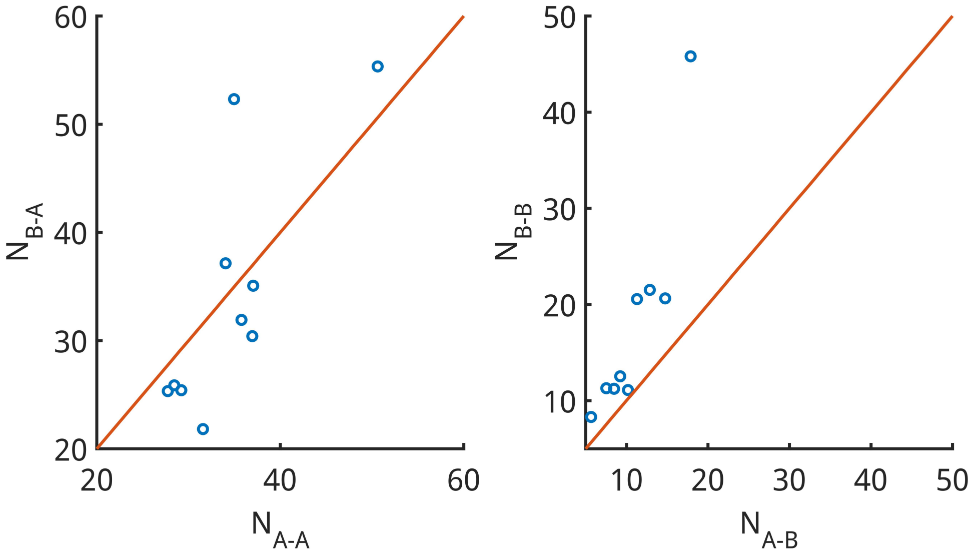


**Figure S21. A robustness test for the association between gene co-regulation and compartmentalization, corresponding to Figure 4A.** The criterion we used here in identifying highly correlated gene pairs is 99.9$^{th}$ percentile (see methods).

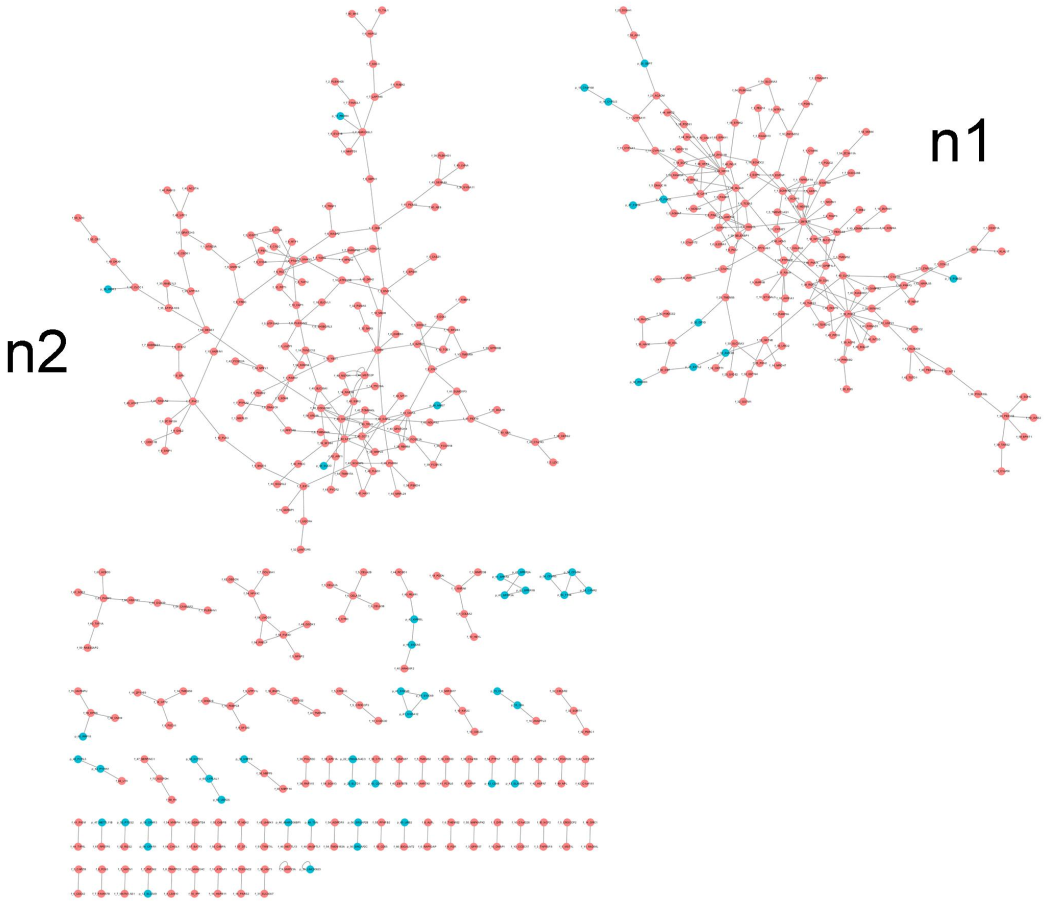

**Figure S22. Gene network in liver.** Red circle, forest gene; blue circle, prairie gene. n1, liver-specific sub-network; n2, the more generic sub-network.

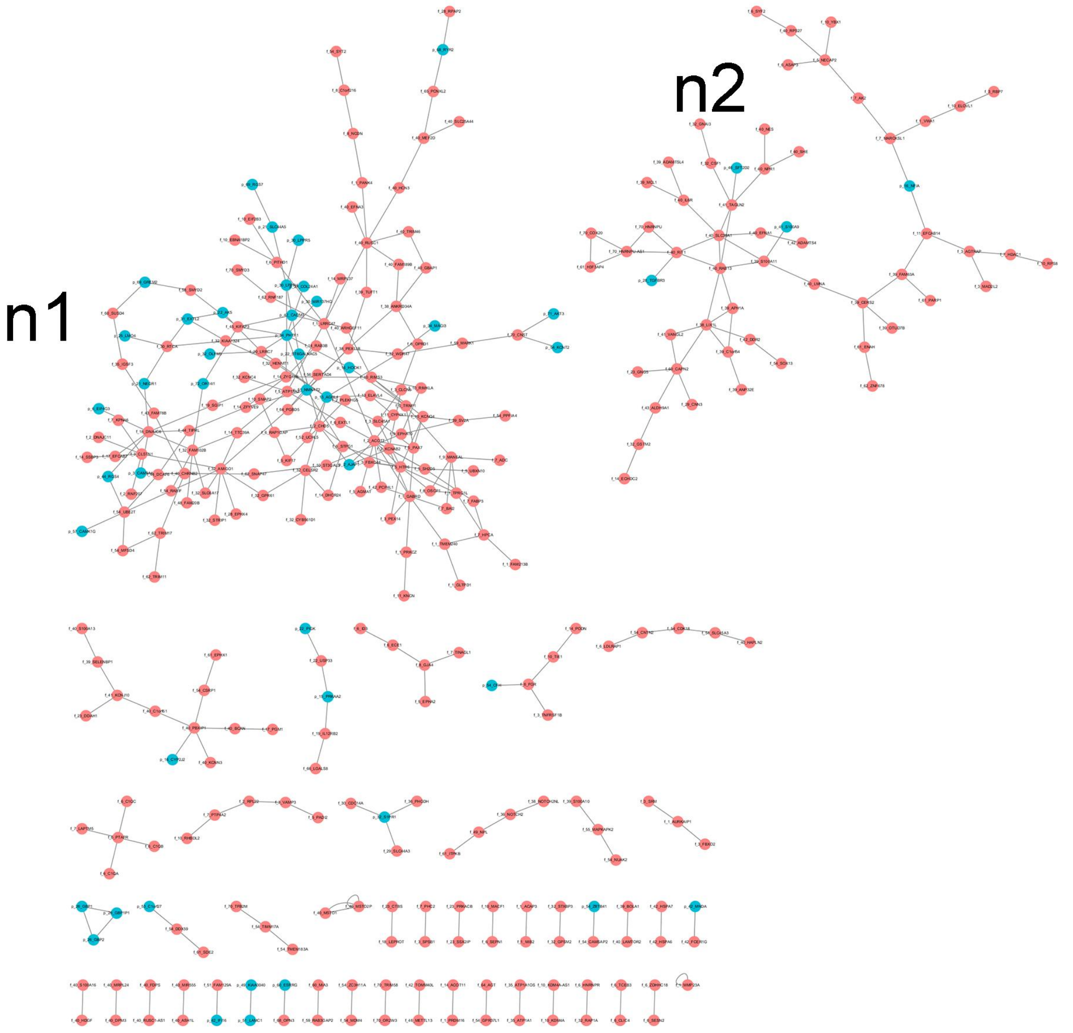


**Figure S23. Gene network in cortex BA9.** Red circle, forest gene; blue circle, prairie gene. n1, cortex-specific sub-network; n2, more generic sub-network.

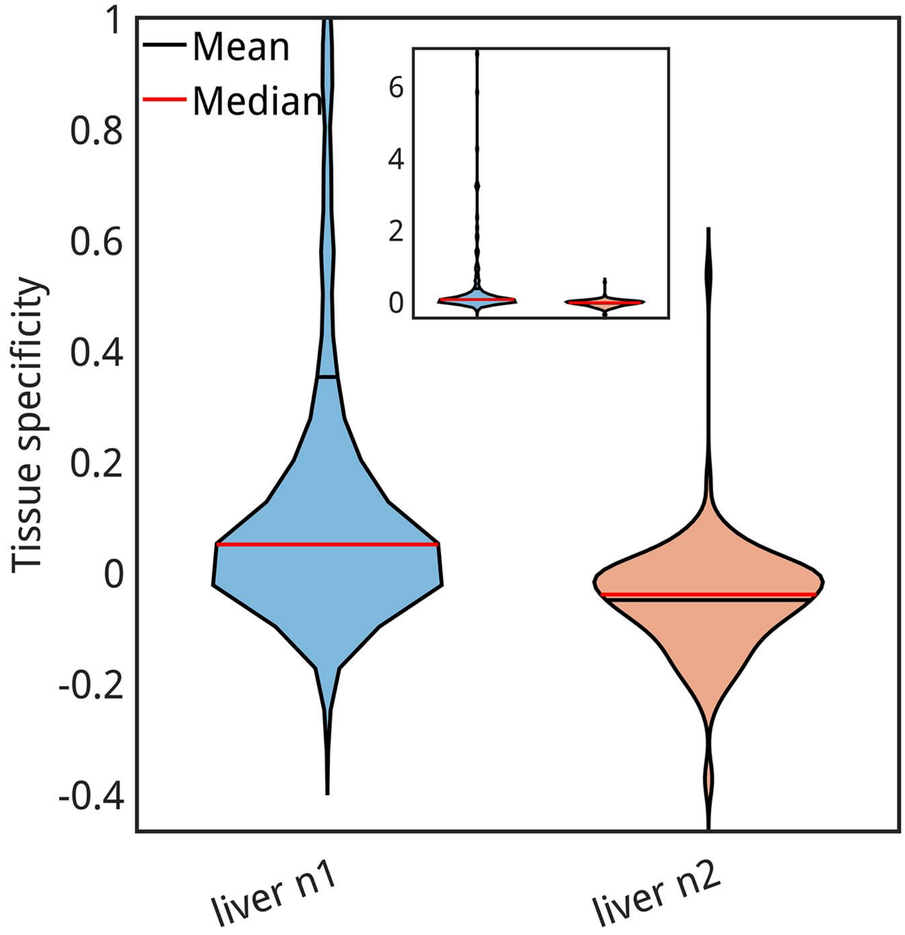


**Figure S24. The tissue specificity distribution of two liver sub-networks.** Inner, the intact and unamplified figure.

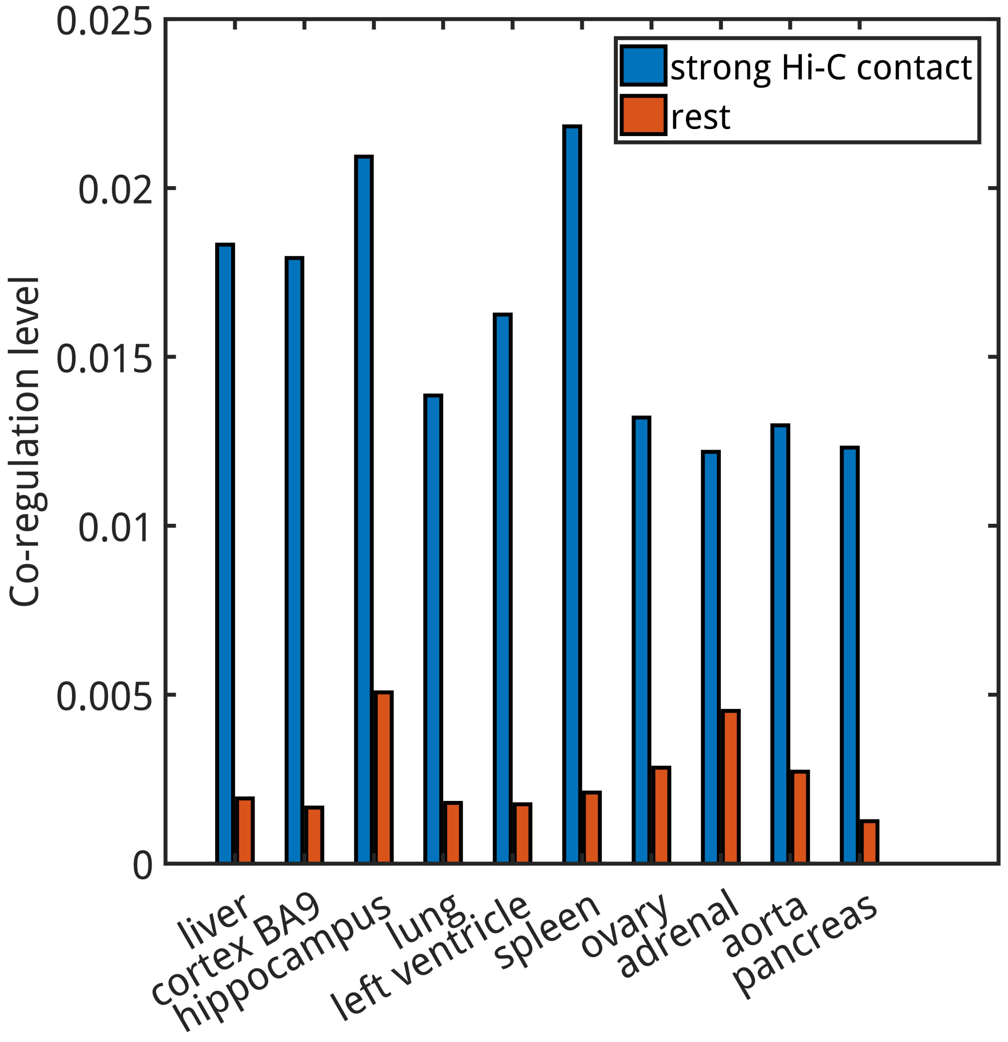


**Figure S25. A robustness test for how spatial contact affects gene co-regulation level, corresponding to Figure 4E.** The criterion we used here in identifying highly contacted gene pairs is 90$^{th}$ percentile (top 10%, see methods) and average values were shown.
