## Supplementary File 1 for "Toward an understanding of the relation between gene regulation and 3D genome organization"

38 tissues

Adipose_subcutaneous
Adipose_visceral
Adrenal_gland
Artery_aorta
Artery_coronary
Artery_tibial
Brain_basal_ganglia
Brain_cerebellum
Brain_other
Breast
Colon_sigmoid
Colon_transverse
Esophagus_mucosa
Esophagus_muscularis
Fibroblast_cell_line
Gastroesophageal_junction
Heart_atrial_appendage
Heart_left_ventricle
Intestine_terminal_ileum
Kidney_cortex
Liver
Lung
Lymphoblastoid_cell_line
Minor_salivary_gland
Ovary
Pancreas
Pituitary
Prostate
Skeletal_muscle
Skin
Spleen
Stomach
Testis
Thyroid
Tibial_nerve
Uterus
Vagina
Whole_blood
